# *Helicobacter pylori* accelerates KRAS-dependent gastric dysplasia

**DOI:** 10.1101/2020.10.06.328518

**Authors:** Valerie P. O’Brien, Amanda Koehne, Julien Dubrulle, Armando E. Rodriguez, Christina K. Leverich, Paul Kong, Jean S. Campbell, Robert H. Pierce, James R. Goldenring, Eunyoung Choi, Nina R. Salama

## Abstract

More than 80% of gastric cancer is attributable to stomach infection with *Helicobacter pylori* (*Hp*), even though the bacterium is not always present at time of diagnosis. Infection is thought to lead to cancer by promoting the accumulation of oncogenic mutations downstream of inflammation; once oncogenic pathways become activated, infection may become dispensable for cancer development. Gastric preneoplastic progression involves sequential changes to the tissue, including loss of parietal cells, spasmolytic polypeptide-expressing metaplasia (SPEM), intestinal metaplasia (IM) and dysplasia. In mice, active KRAS expression recapitulates these tissue changes in the absence of *Hp* infection. This model provides an experimental system to investigate whether *Hp* infection has additional roles in preneoplastic progression, beyond initiating inflammation. Mice were assessed by evaluating tissue histology, gene expression changes, the immune cell repertoire, and expression of metaplasia and dysplasia markers. Compared to *Hp*-/KRAS+ mice, *Hp*+/KRAS+ mice had i) severe T cell infiltration and altered macrophage polarization; ii) altered expression of metaplasia markers, including increased expression of CD44v9 (SPEM) and decreased expression of TFF3 (IM); iii) more dysplastic (TROP2+) glands; and iv) greater proliferation of metaplastic and dysplastic glands. *Hp* was able to persistently colonize the stomach during the onset of these tissue changes, and eradication of *Hp* with antibiotics prevented metaplastic, dysplastic and proliferation marker changes. Collectively, these results suggest that gastric preneoplastic progression differs between *Hp*+ and *Hp*-cases, and that sustained *Hp* infection can promote the later stages of gastric preneoplastic progression, in addition to its established role in initiating chronic inflammation.

## Introduction

About 13% of the global cancer burden in 2018 was attributable to carcinogenic infections ^1^, and *Helicobacter pylori* (*Hp*)-associated gastric cancer accounted for the largest proportion of these cancers ^2^. More than 77% of new gastric cancer cases, and more than 89% of new non-cardia gastric cancer cases, were attributable to infection with *Hp* ^1^, a bacterium that colonizes the stomach of half the world’s population ^3^. However, *Hp* infection confers only a 1 to 2% lifetime risk of developing stomach cancer ^4^ and thus a complex interplay between the bacterium and host is presumed to lead to cancer development in only some individuals.

The exact mechanisms through which *Hp* infection promotes gastric cancer remain largely elusive. *Hp* infection typically occurs during childhood and always causes chronic inflammation (gastritis) ^5^. *Hp*-dependent chronic inflammation promotes the accumulation of reactive oxygen species and other toxic products that cause mutations in gastric epithelial cells ^6-8^. Early studies using tissue histology rarely detected *Hp* in tumors, leading to a belief that *Hp* triggers the initial inflammatory insult in the stomach, but that *Hp* is essentially irrelevant by the time gastric cancer is detected; in other words, once chronic gastric inflammation develops and oncogenic pathways are activated, the presence of *Hp* is no longer necessary to promote metaplastic changes that lead to cancer. However, more sensitive molecular methods detect *Hp* in about half of tumors ^9-11^, and eradication of *Hp* combined with tumor resection helps prevent tumor recurrence ^12^, suggesting that *Hp* may promote the later stages of metaplasia and cancer development in at least some individuals.

Beyond eliciting oncogenic mutations, the mechanism(s) through which chronic gastritis might promote gastric cancer development is not well understood ^13^. Humans generally develop a strong T_h_1 and T_h_17 immune response against *Hp* that helps control the infection ^14-16^. This T cell response does not clear the infection and furthermore can drive immunopathology in the gastric mucosa ^17, 18^, and *Hp* infection can disrupt normal T cell function through multiple mechanisms ^13, 19, 20^. Thus, T cells can play both protective and detrimental roles during *Hp* stomach infection. More broadly, anticancer immunity in the context of gastric cancer is not well understood. A better understanding of how active *Hp* infection may impact gastric inflammation in the context of metaplasia and cancer development may lead to the discovery of new drug targets or therapeutic strategies.

The *Mist1-Kras* mouse is one of the only existing mouse models to recapitulate the progression from healthy gastric epithelium to spasmolytic polypeptide-expressing metaplasia (SPEM), intestinal metaplasia (IM) and dysplasia ^21^. This model utilizes KRAS, a GTPase signaling protein of the Ras (Rat Sarcoma) family that regulates cell survival, proliferation and differentiation ^22, 23^. Molecular profiling studies have shown that about 40% of gastric tumors have signatures of RAS activity ^24, 25^. In the mouse model, treatment with tamoxifen (TMX) induces the expression of a constitutively active *Kras* allele (G12D) in the gastric chief cells. Within one month, SPEM develops in 95% of corpus glands, and over the next three months progresses to IM ^21^. Thus, active KRAS expression in mice serves as a tool to recapitulate changes that, in humans, are induced by years of inflammation due to *Hp* infection. We used *Mist1-Kras* mice to test our hypothesis that *Hp*, if present during metaplasia and dysplasia, could impact pathology. We found that sustained *Hp* infection coupled with active *KRAS* expression led to severe inflammation, altered metaplasia marker expression, and increased cell proliferation and dysplasia compared to *Hp*-/KRAS+ mice. Thus, the course of gastric neoplastic progression may differ depending on whether *Hp* is present during the later stages of disease progression.

## Results

### *Hp* infection worsens gastric immunopathology in mice expressing active KRAS

To assess whether *Hp* impacts KRAS-driven metaplasia, we performed concomitant infection/induction experiments in *Mist1-Kras* mice. First mice were infected with *Hp*, or mock-infected, and the next day mice were treated with tamoxifen (TMX) to induce active KRAS expression in stomach chief cells, or sham-induced. After two, six or 12 weeks, mice were humanely euthanized and stomachs were aseptically harvested and used for downstream analyses (**Figure 1**). Formalin-fixed, paraffin-embedded tissue sections were used for histological analysis of the corpus (**Figure 2**), where active KRAS is expressed in TMX-induced *Mist1-Kras* mice. Compared to *Hp*-/KRAS-mice (**Figure 2A and B**), *Hp* infection alone caused modest inflammation at two weeks that increased over time, with loss of parietal cells by six weeks and moderate surface epithelial hyperplasia by 12 weeks (**Figure 2C and D**). Mice expressing active KRAS had far more striking changes to the tissue over time (**Figure 2E-H**). To quantify the effects of *Hp* infection in this model, a blinded analysis was performed to assess inflammation, oxyntic atrophy (loss of parietal cells), and surface epithelial hyperplasia in active KRAS-expressing mice (**Figure 3A-C**).

**Figure 1.**
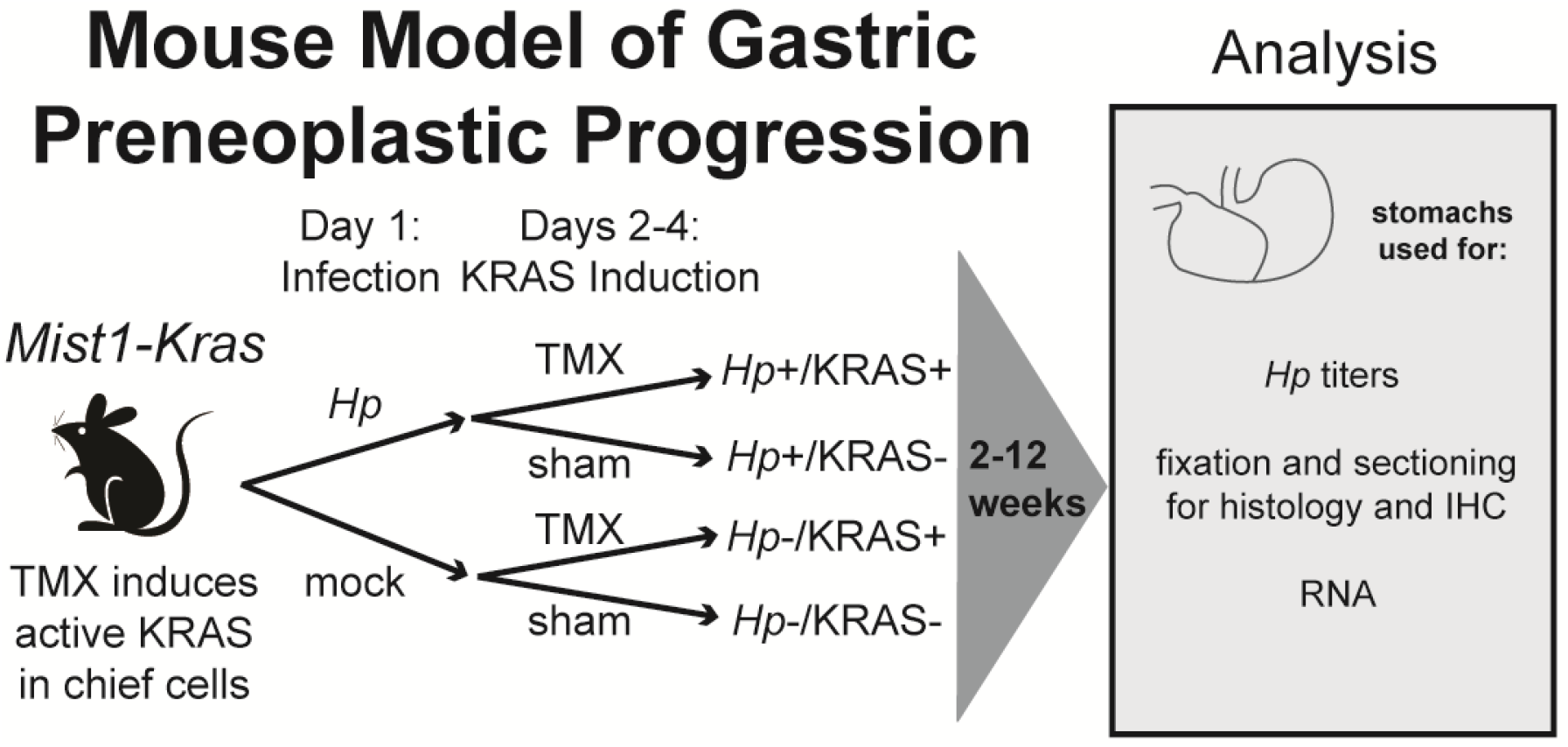
*Mist1-Kras* mice were used to assess whether and how *Hp* infection alters gastric preneoplastic progression. On day one, mice are infected with *H. pylori* (*Hp*) by oral gavage, or mock-infected. On days two through four, mice receive daily injections with tamoxifen (TMX) to induce a constitutively active *Kras* allele (G12D) in the chief cells (*Mist1*-expressing) of the stomach. After two, six or 12 weeks, mice are humanely euthanized and the glandular stomach (excluding forestomach region) is assessed as indicated.

**Figure 2.**
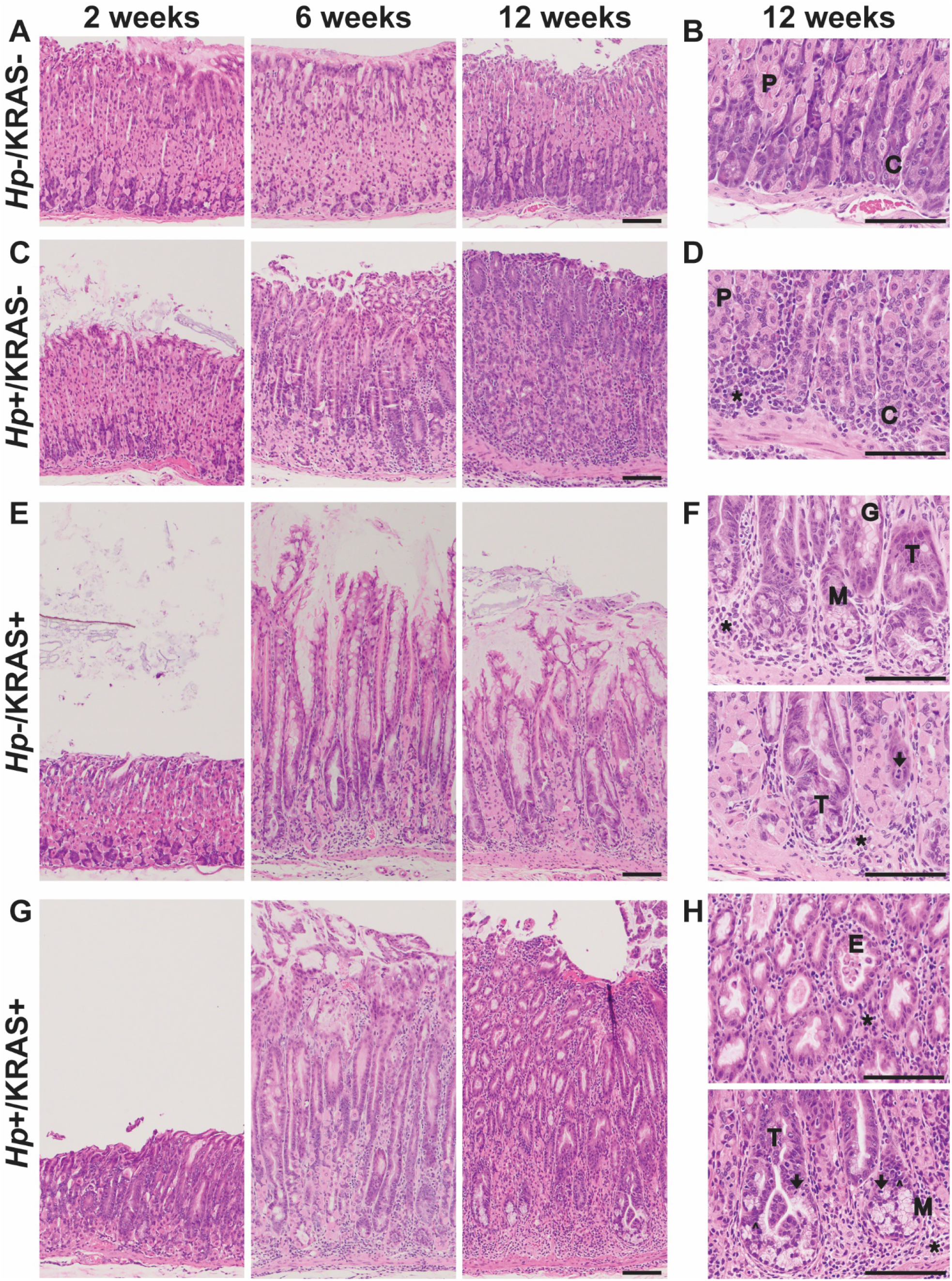
Concomitant *Hp* infection and active KRAS expression changes tissue histology. Shown are representative images of corpus tissue from formalin-fixed, paraffin-embedded, hematoxylin & eosin-stained sections from stomachs obtained from *Hp*-/KRAS-(**A and B**), *Hp*+/KRAS-(**C and D**), *Hp*-/KRAS+ (**E and F**), and *Hp*+/KRAS+ (**G and H**) mice (n=10-16 per group) after two, six or 12 weeks as indicated. Scale bars, 100 µm. Examples of the following morphological features are designated in the higher magnification images in **B, D, F** and **H**: chief cells (C); glandular ectasia (E); goblet-like cell morphology (G); hyperchromatic nuclei (carets); immune cells (asterisks); mitotic figures (arrows); mucus (M); parietal cells (P); tortuous glands (T).

**Figure 3.**
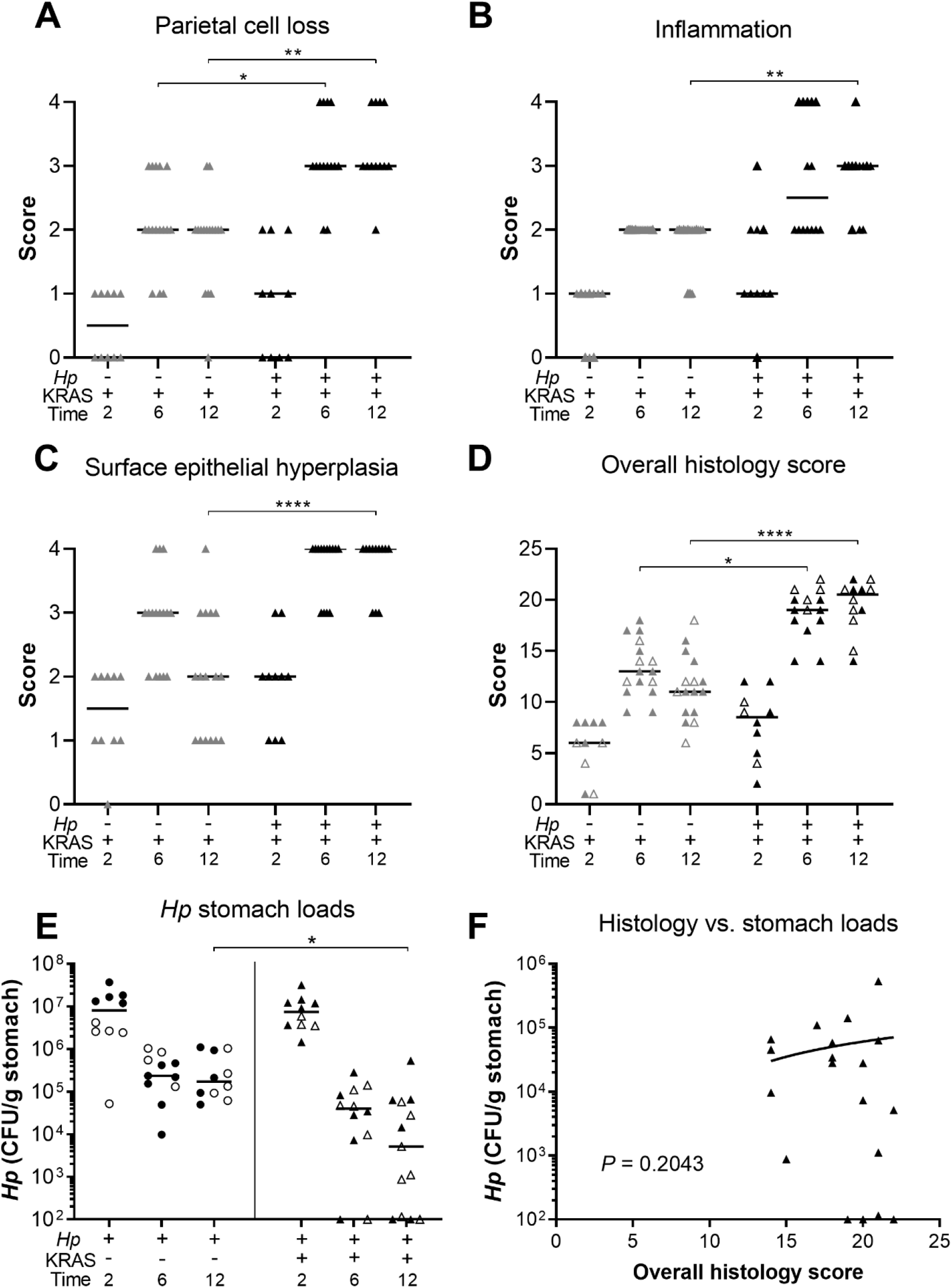
*Hp*+/KRAS+ mice have severe gastric immunopathology marked by inflammation, loss of parietal cells and surface epithelial hyperplasia. Stomachs from n=10-16 mice per group were evaluated for tissue pathology and bacterial colonization. (**A-D**) Hematoxylin-and-eosin-stained corpus tissue was assessed for parietal cell loss (oxyntic atrophy) (**A**), inflammation (**B**), surface epithelial hyperplasia (**C**) and the composite histological activity index (**D**) in a blinded fashion using the Rogers criteria ^26^. (**E**) *Hp* loads were assessed by quantitative culture; mice with no detectable colonization were plotted at the limit of detection. (**F**) Comparison of *Hp* loads and overall histology score (from **D**) from the same mouse at six or 12 weeks. Significance was assessed by Spearman correlation. Data are combined from N=2 independent mouse experiments per time point. Data points represent actual values for each individual mouse and bars indicate median values. Statistically significant comparisons are indicated by: * *P* < 0.05, ** *P* < 0.01, **** *P* < 0.0001, Kruskal-Wallis test with Dunn’s multiple test correction. CFU/g, colony-forming units per gram stomach tissue. In **D** and **E**, open vs. closed symbols distinguish between biological replicates.

KRAS expression caused changes to the corpus epithelium that were apparent within two weeks, with a moderate degree of inflammation, surface epithelial hyperplasia, and some loss of parietal cells. These changes were slightly more severe in a subset of *Hp*+/KRAS+ mice, but overall there were no significant histological differences between *Hp*-/KRAS+ and *Hp*+KRAS+ mice at this early time point (**Figure 2E and G**). By six weeks, each of these parameters became more severe, and notably, parietal cell loss was significantly greater in *Hp*+/KRAS+ mice compared to *Hp*-/KRAS+ mice (**Figure 3A**). By 12 weeks, *Hp*-/KRAS+ mice had mucinous cells, in line with previous observations ^21^ (**Figure 2F**). *Hp*+KRAS+ mice looked different from *Hp*-/KRAS+ mice, with loss of normal basal polarity of epithelial cells, and gland architecture that was severely disrupted, including forked or star-shaped gland structure indicative of extensive branching and disorganized maturation (**Figure 2G**). As well, these mice had hyperchromatic nuclei with variations in nuclear size, which can indicate dysplasia. Moreover, at 12 weeks *Hp*+/KRAS+ mice had significantly increased inflammation, parietal cell loss and surface epithelial hyperplasia compared to *Hp*-/KRAS+ mice (**Figure 3A-C**). Finally, the overall histology score (histological activity index), which sums the above scores along with scores for other parameters like epithelial defects and hyalinosis (**Supplementary Figure S1**) and which thus indicates the degree of overall immunopathology ^26^, was significantly increased in *Hp*+/KRAS+ mice compared to *Hp*-/KRAS+ mice at both six and 12 weeks (**Figure 3D**). Thus, concomitant *Hp* infection and active KRAS expression in the corpus leads to histopathological changes to the tissue within six weeks that become more severe by 12 weeks.

The striking inflammation seen in *Hp*+/KRAS+ mice compared to *Hp*+/KRAS-mice might be expected to eliminate *Hp* infection. However, *Hp* was recovered from most KRAS+ mice by stomach culturing (**Figure 3E**), demonstrating that the bacterium could to some extent withstand the severe inflammation of the preneoplastic stomach. At two weeks, *Hp* titers were not significantly different between *Hp*+/KRAS- and *Hp*+/KRAS+ mice, suggesting that the early histopathological changes did not impact bacterial colonization. In sham-induced (KRAS-) mice, *Hp* titers were similar at six and 12 weeks, and in both cases were lower than at two weeks, likely due to the onset of adaptive immunity to control the infection. However, in *Hp*+/KRAS+ mice, the contraction of the *Hp* population was greater, with *Hp* recovered from only ten out of 12 animals at six weeks and ten out of 13 animals at 12 weeks. *Hp* could be detected within glands by immunofluorescence microscopy (**Supplementary Figure S2**). No differences in titer or overall histology score were observed between male and female mice. Interestingly, stomach *Hp* loads were not correlated with histology scores (**Figure 3F**). We therefore hypothesized that the host inflammatory response to *Hp* infection might contribute to *Hp*-dependent tissue changes.

### *Hp* infection increases and alters KRAS-driven inflammation

Activated KRAS expression itself elicits inflammation: by 12 weeks, the median corpus inflammation score in *Hp*-/KRAS+ mice was two (**Figure 3B**), denoting coalescing aggregates of inflammatory cells in the submucosa and/or mucosa ^26^. In this histological evaluation, the addition of *Hp* significantly increased inflammation by 12 weeks (*P* < 0.01), with the median score rising to three (denoting organized immune cell nodules in the submucosa and/or mucosa). To characterize the nature of the inflammation, gene expression changes at 12 weeks were assessed with a NanoString mouse immunology panel (**Figure 4 and Supplementary Table S2**). Transcripts of *Cd45*, a pan-immune cell marker, were significantly increased in *Hp*+/KRAS-mice and especially in *Hp*+/KRAS+ mice compared to *Hp*-mice (**Supplementary Figure S3**), suggesting greater numbers of immune cells in infected mice. Of the 561 genes in the panel, 60 had no detectable expression among any mice (n=25) and were excluded from subsequent analysis. Hierarchical clustering was performed on the remaining 501 genes and revealed distinct clustering of the treatment groups (**Figure 4A**). All *Hp*+ mice clustered separately from *Hp*-mice, demonstrating that infection had the greatest impact on gene expression. Within both the *Hp*+ and the *Hp*-cluster, KRAS+ mice clustered separately from KRAS-mice, suggesting that active KRAS expression also impacted inflammatory gene expression, though to a lesser extent than *Hp* infection did.

**Figure 4.**
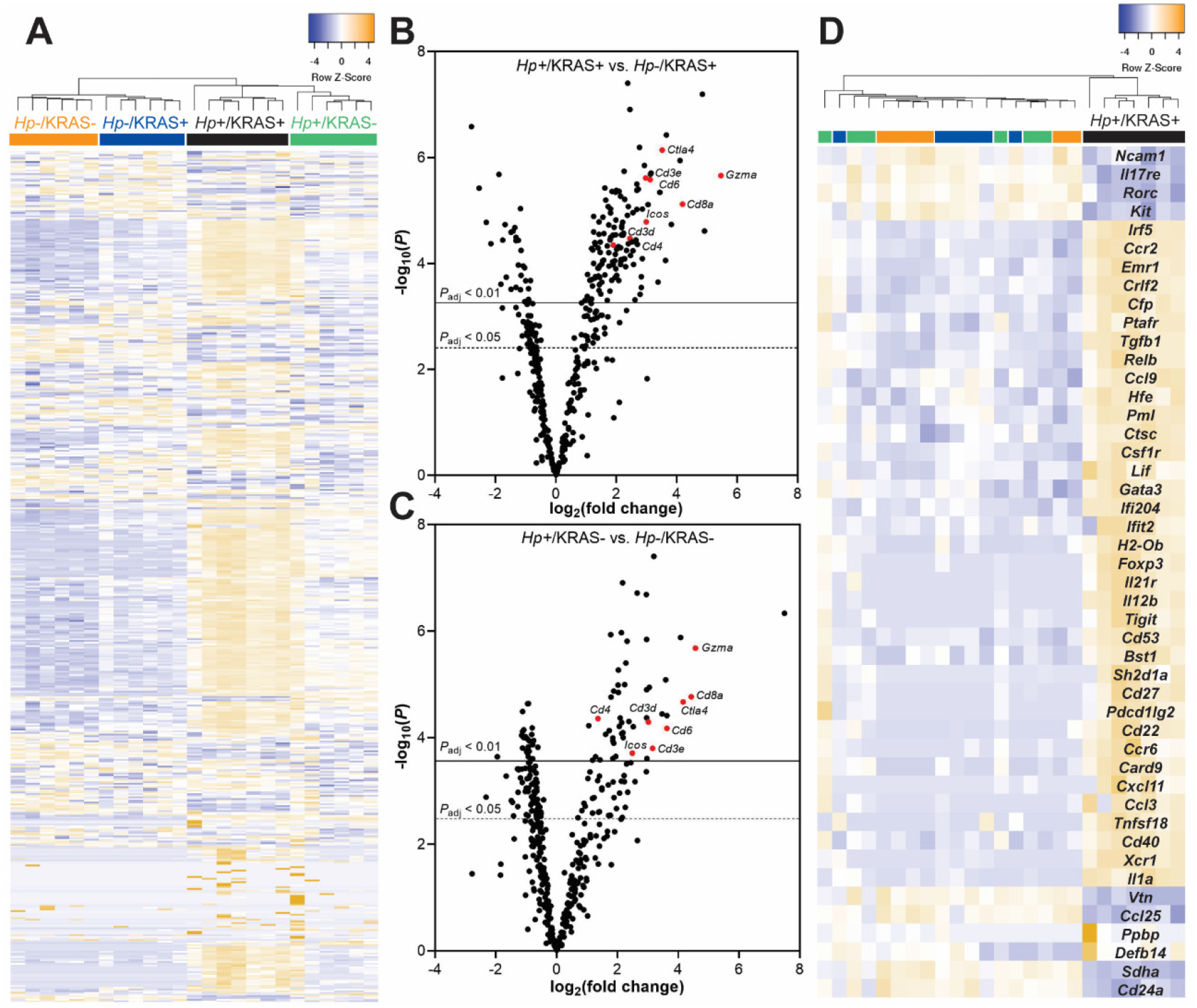
A unique inflammatory gene signature exists in *Hp*+/KRAS+ mice at 12 weeks. RNA was extracted from stomach sections from *Hp*+/-, KRAS+/- mice at 12 weeks and immune-related gene expression was detected with the NanoString nCounter Mouse Immunology Panel. (**A**) Expression of 501 genes is shown. Colored bars denote different treatment groups. (**B and C**) Volcano plots show the fold change and *P* values of all genes in the panel, for *Hp*+/KRAS+ mice vs. *Hp*-/KRAS+ mice (**B**) and *Hp*+/KRAS-mice vs. *Hp*-/KRAS-mice (**C**). A subset of T cell-related genes is shown in red and labeled. Lines show genes meeting the threshold for significance after correction with the Benjamini-Yekutieli procedure. (**D**) Expression of 46 genes (see text) that were uniquely differentially expressed in *Hp*+/KRAS+ mice vs. all other groups is shown. The dendrograms at the top of the heat maps were produced by hierarchical clustering of gene expression. Data comes from N=1 NanoString experiment with n=6-7 mice per group from N=2 independent mouse experiments.

Next we assessed gene expression patterns in the different mouse groups. Compared to *Hp*-/KRAS+ mice, *Hp*+/KRAS+ mice had 235 significantly differentially expressed genes (DEGs) (*P*_adjusted_ < 0.05) (**Figure 4B**). Several of the most highly upregulated genes, including *Cd3d, Cd3e, Cd4, Cd8a, Gzma, Ctla4, Icos* and *Cd6*, implicated a strong T cell response, in accordance with previous studies in humans and naive animal models ^27, 28^. Likewise, compared to *Hp*-/KRAS-mice, *Hp*+/KRAS-mice had 177 DEGs, with *Cd3d, Cd3e, Cd4, Cd8a, Gzma, Ctla4, Icos* and *Cd6* once again highly significantly differentially expressed (**Figure 4C**). Thus, many of the gene expression differences seen in *Hp*+/KRAS+ mice vs. *Hp*-/KRAS+ mice are likely reflective of a general pattern of *Hp*-mediated inflammation that is independent of the metaplastic state of the tissue. However, we identified a unique inflammatory gene signature in *Hp*+/KRAS+ mice (**Figure 4D**), demonstrating that the inflammation observed in this group is not only of a greater magnitude than in the other groups, but also of a different nature. We identified 46 genes whose expression (normalized to *Hp*-/KRAS-mice) was >2-fold increased or decreased in *Hp*+/KRAS+ mice, but <1.5-fold increased or decreased in *Hp*+/KRAS- and *Hp*-/KRAS+ mice. Many of these genes implicated T cells (*Ccr6, Cd27, Cd53, Cxcl11, Foxp3, Gata3, Il12b, Pdcd1lg2* [PD-L2], *Tigit*, and *Tnfsf18* upregulated; *Il17re* downregulated) and macrophages (*Ccl3, Csf1r, Emr1* [F4/80], *Il1a* and *Irf5* upregulated). As well, most markers of T cell exhaustion ^29, 30^ were only strongly expressed in *Hp*+/KRAS+ mice (**Supplementary Figure S4**). Thus, even though both *Hp*+/KRAS+ mice and *Hp*+/KRAS-mice had significant upregulation of T cell-related genes compared to their *Hp*-counterparts, the addition of active KRAS may impact the nature of T cell polarization and function.

In animal models, immune pressure due to chronic *Hp* infection results in loss of function of the *Hp* type IV secretion system (T4SS) ^31^. *Hp* strains isolated from long-term experimental infections of C57BL/6 mice (but not *Rag1* mice deficient in adaptive immune responses), gerbils and monkeys lose their ability to elicit IL-8 secretion by gastric epithelial cells *in vitro* ^31^. In line with these observations, we found that approximately 50% of *Hp* strains isolated from 12+ week infections of KRAS-mice had lost their T4SS activity (**Supplementary Figure S5**). Surprisingly, *Hp* strains isolated from KRAS+ mice were no more likely to lose their T4SS activity, despite the severe inflammation seen in these animals.

### *Hp*+/KRAS+ mice have T cells throughout the lamina propria and fewer M2 macrophages

To detect immune cell subsets in the corpus of *Hp*-/KRAS+ vs. *Hp*+/KRAS+ mice at 12 weeks, we performed multiplex fluorescent immunohistochemistry (IHC) with the following markers: for T cells, CD3, CD4, CD8α, FOXP3 (regulatory T cell marker) and PD-1 (T cell exhaustion marker); for macrophages, F4/80 and the polarization markers MHC class II (M1 macrophages) and CD163 (M2 macrophages) (**Figure 5**). HALO software was used to detect and enumerate immune cell subsets (**Supplementary Figure S6**). In *Hp*-/KRAS+ mice we detected moderate numbers of CD3+ T cells, most of which were CD4+, and a few of which were CD8α+ (**Figure 5A and Supplementary Figure S6A**). *Hp*+/KRAS+ mice had significantly more CD3+ T cells, but the proportion of CD4+ vs. CD8α+ cells was similar, with more CD4+ than CD8α+ cells. Interestingly, most of the CD3+ cells in *Hp*-/KRAS+ mice expressed FOXP3 and PD-1 (**Figure 5B**), suggesting they may be activated regulatory T cells ^32^. In *Hp*+/KRAS+ mice, there were significantly more FOXP3+ cells (**Supplementary Figure S6A**), some of which were PD-1 double-positive (**Figure 5B**). However, many CD3+ cells did not express either of these markers, suggesting they may be different T cell subsets than are found in *Hp*-/KRAS+ mice, and/or NK cells. Cell localization was also different between treatment groups: in *Hp*-/KRAS+ mice, most T cells were located at the base of the glands, whereas in *Hp*+/KRAS+ mice, T cells were located throughout the glands. Finally, both groups of mice had F4/80+ cells throughout the lamina propria (**Figure 5C and Supplementary Figure S6B**), suggesting presence of macrophages or eosinophils ^33^. We previously found that M2 macrophages promoted SPEM progression in mice and were associated with human SPEM and IM ^34^. In *Hp*-/KRAS+ mice, some F4/80+ cells were dual-positive for the M2 polarization marker CD163, in line with previous findings ^21^, and some were dual-positive for the M1 polarization marker MHC class II. *Hp*+/KRAS+ mice had similar numbers of F4/80+/MHC class II+ cells present, but significantly fewer F4/80+/CD163+ cells (**Supplementary Figure S6B**), suggesting altered macrophage polarization; most CD163 signal was observed in the gland lumen, likely non-specific staining due to mucus binding. These IHC experiments confirm our gene expression-based findings that inflammation in *Hp*+/KRAS+ mice is not only more severe than in *Hp*-/KRAS+ mice, but is also altered in nature.

**Figure 5.**
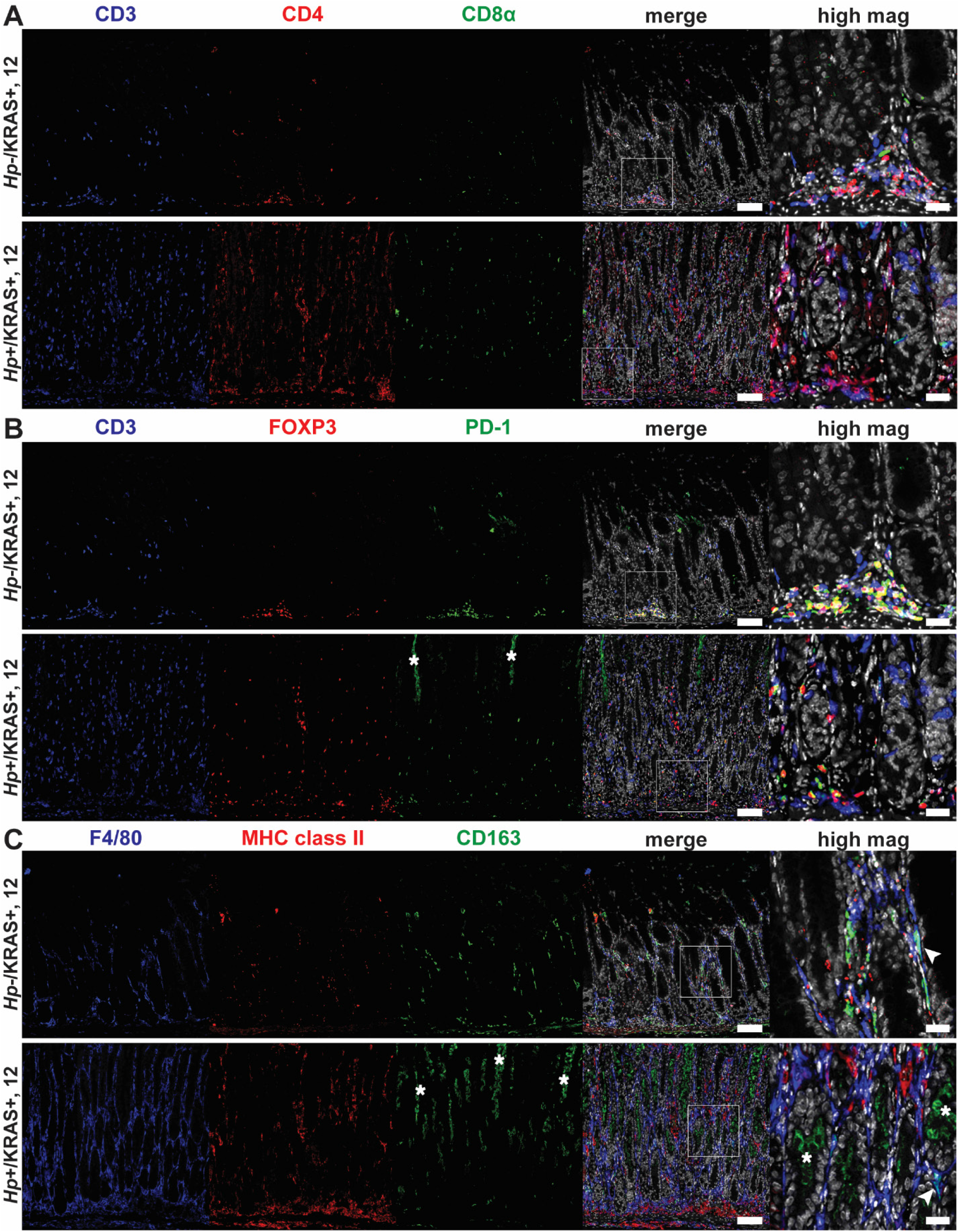
Multiplex immunohistochemistry demonstrates more T cells and fewer M2 macrophages in *Hp*+/KRAS+ mice. Corpus tissue from *Hp*-/KRAS+ and *Hp*+/KRAS+ mice obtained after 12 weeks was assessed by multiplex immunohistochemistry and representative images are shown. White boxes denote regions shown at higher magnification. (**A**) CD3 suggests T cells, CD4 indicates helper T cells, CD8α indicates cytotoxic T cells. (**B**) CD3 suggests T cells, FOXP3 indicates regulatory T cells, PD-1 is a marker of exhausted T cells. (**C**) F4/80 suggests macrophages, MHC class II indicates M1 macrophages, CD163 indicates M2 macrophages, arrowhead indicates an F4/80+/CD163+ cell. Scale bars for low magnification (“merge”) images, 100 µm; for high magnification images, 25 µm. Data are from N=1 multiplex immunohistochemistry experiment, with n=7 mice per group from N=2 independent mouse experiments. Asterisks denote examples of non-specific staining as determined by luminal location, lack of DAPI signal and lack of co-localization with relevant markers (**B**, CD3 and **C**, F4/80).

### *Hp* infection alters metaplasia marker expression

We wondered whether the changes in tissue histology observed in *Hp*+/KRAS+ mice (**Figure 2**) reflected changes to the nature of metaplasia in these mice. To detect differences in hyperplasia, metaplasia and cell proliferation in corpus tissue from *Hp*+/KRAS+ vs. *Hp*-/KRAS+ mice over time (**Figure 6A-C**), we used: conjugated lectin from *Ulex europaeus* (UEA-I), which binds alpha-L-fucose, to detect foveolar (pit cell) hyperplasia; conjugated *Griffonia simplicifolia* lectin II (GS-II), which binds α- or β-linked N-acetyl-D-glucosamine, to detect mucous neck cells and SPEM cells; anti-CD44v10 (orthologous to human CD44v9, referred to herein as “CD44v”) to detect SPEM cells ^35^; anti-TFF3 and anti-MUC2 to detect IM (goblet) cells ^36^ (verified by staining of mouse intestine as shown in **Supplementary Figure S7**); and anti-KI-67 to detect proliferating cells. We assessed differences through quantitative and semi-quantitative analysis of three to five images per mouse (**Figure 6D-H**). No difference was observed in UEA-I staining among the treatment groups (**Supplementary Figure S8**), suggesting that *Hp* infection did not impact foveolar hyperplasia development in this model. In *Hp*-/KRAS+ mice, GS-II staining was observed at the base of the glands at six weeks, co-localizing with CD44v, demonstrating SPEM (**Figure 6A, D, E**). As well, most mice had robust, cell-associated TFF3 staining (**Figure 6A and F**) and a low degree of MUC2 staining (**Figure 6B and G**), suggestive of early IM. At 12 weeks, CD44v and GS-II staining was reduced, TFF3 staining remained robust, and the percentage of MUC2+ glands increased, suggesting a transition from SPEM to IM in these mice, consistent with previous findings ^21^. In *Hp*+/KRAS+ mice, GS-II and MUC2 staining patterns were similar, with GS-II decreasing and MUC2 increasing between six and 12 weeks (**Figure 6A, B, D, G**). However, CD44v and TFF3 exhibited a different pattern. At six weeks, *Hp*+/KRAS+ mice had greater CD44v staining and less TFF3 staining compared to *Hp*-/KRAS+ mice (**Figure 6A, E, F**). By 12 weeks, CD44v staining waned somewhat but remained higher than in *Hp*-/KRAS+ mice, and TFF3 staining further diminished compared to *Hp*-/KRAS+ mice. Taken together, these results demonstrate that *Hp* infection alters the kinetics of metaplasia development in KRAS+ mice. Metaplasia marker expression was not detected in KRAS-mice (**Supplementary Figure S9**). Of note, no differences in immunostaining patterns or quantification were observed in KRAS+ mice at two weeks (**Supplementary Figure S10**), suggesting that *Hp*-driven metaplastic changes take longer than two weeks to become apparent.

**Figure 6.**
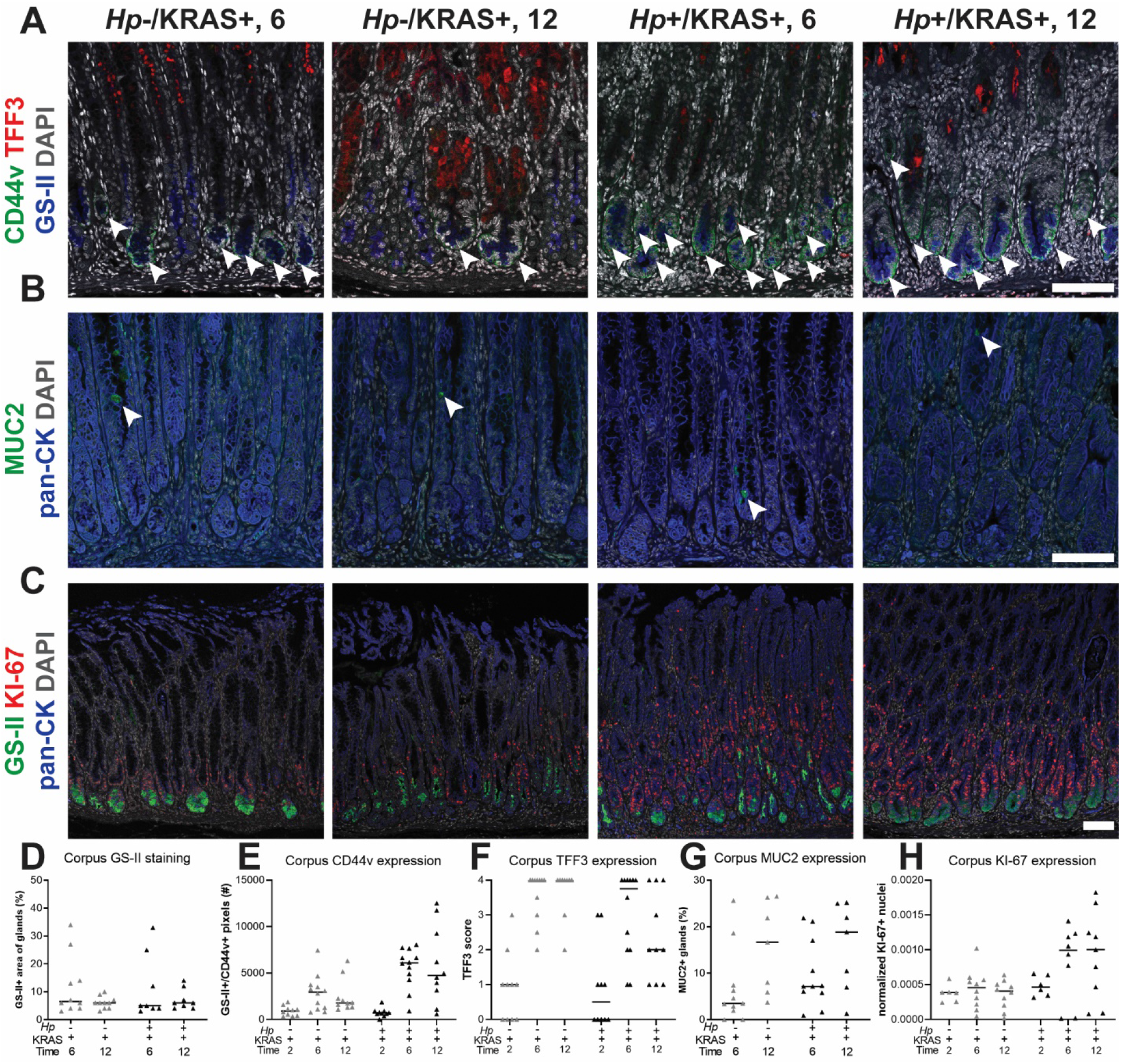
The kinetics and molecular nature of metaplasia development are altered in *Hp*+/KRAS+ mice. Corpus tissue from *Hp*-/KRAS+ and *Hp*+/KRAS+ mice obtained after two, six or 12 weeks (N=2 independent mouse experiments per time point and n=6-12 mice per group) was assessed for metaplasia via immunofluorescence microscopy. (**A-C**) Representative images are shown and scale bars are 100 µm. (**A**) Stomachs were stained with antibodies against CD44v (green; arrowheads) and TFF3 (red), the lectin GS-II (blue) and DAPI (grey) in N=3 staining experiments. (**B**) Stomachs were stained with antibodies against MUC2 (green; arrowheads) and pan-cytokeratin (blue) and DAPI (grey) in N=2 staining experiments. (**C**) Stomachs were stained with antibodies against KI-67 (red) and pan-cytokeratin (blue), the lectin GS-II (green) and DAPI (grey) in N=3 experiments. (**D-H**) Three to five representative images per mouse were quantitatively or semi-quantitatively assessed and the median value for each mouse is plotted. Bars on the graphs indicate the median value for each mouse group. (**D**) The percentage of cytokeratin-positive epithelial tissue that was dual-positive for GS-II staining was detected. (**E**) The number of GS-II+/CD44v+ pixels per image was quantified. (**F**) TFF3 staining was semi-quantitatively scored in a blinded fashion. Non-specific staining was not included in the score (see **Figure S9**). (**G**) The percentage of MUC2+ glands was determined by counting. (**H**) KI-67+/DAPI+ nuclei were enumerated and normalized to the DAPI content (total number of DAPI+ pixels) of each image.

We observed mitotic figures in KRAS+ mice at 12 weeks (**Figure 2F and H**), suggesting increased cell division. Previously, patients with intestinal metaplasia were found to have significantly increased cellular proliferation (assessed by KI-67 staining) in biopsy tissue compared to healthy controls and patients with chronic active gastritis ^37^. Here we found substantially more KI-67+ nuclei in corpus tissue of *Hp*+/KRAS+ mice than *Hp*-/KRAS+ mice at both six and 12 weeks (**Figure 6C and H**). Most KI-67+ cells were found within the glandular epithelial compartment, not in the lamina propria, and interestingly, the localization of KI-67+ cells was altered in KRAS+ mice. In KRAS-mice, proliferating cells were found in the middle of the glands, where gastric stem cells are found (**Supplementary Figure S9B**). We observed that KI-67+ cells localized toward the base of the glands in *Hp*-/KRAS+ mice (**Figure 6C**). In *Hp*+/KRAS+ mice, KI-67 cells were abundant toward the base of the glands and higher up into the middle of the glands. In both groups of mice, some KI-67+ nuclei were found in GS-II+ cells at the base of the glands, suggesting proliferation of SPEM cells. However, in *Hp*+/KRAS+ mice most KI-67+ nuclei were found above GS-II+ cells, suggesting proliferation of additional cell types beyond those with a SPEM phenotype.

### *Hp* infection increases dysplasia and cancer-associated gene expression

Overexpression of the calcium signal tranducer TROP2 has been implicated in a variety of cancers ^38^, including gastric cancer, where it is associated with worse outcomes ^39^. Notably, TROP2 expression was recently identified as a strong indicator of the transition from incomplete IM to gastric dysplasia in *Mist1-Kras* mice and in human samples ^40^. We observed TROP2+ corpus glands by immunofluorescence microscopy at six and 12 weeks (**Figure 7A and Supplementary Figure S11**). Quantitation using collagen VI as a gland segmentation marker revealed that *Hp*-/KRAS+ mice had TROP2 expression in 0 to 3.6% of glands at six weeks, and 0 to 4.9% of glands at 12 weeks (**Figure 7B and Supplementary Figure S11**). *Hp*+/KRAS+ mice had similar TROP2 expression at six weeks (0.5 to 2.5% of glands), but at 12 weeks had significantly more TROP2+ glands (1.5 to 9.1%, *P* < 0.05). Thus, the addition of *Hp* significantly increased the percentage of TROP2+ glands in the corpus at 12 weeks. In all mice, most regions of TROP2 staining co-localized with KI-67 staining, suggesting proliferation of dysplastic glands. Only a few TROP2+ regions did not harbor KI-67+ cells (**Figure 7A**, asterisk). However, the association between TROP2 and Ki67 was greatest in *Hp*+/KRAS+ mice at 12 weeks, where a median of 86% of TROP2+ glands or gland fragments were KI-67+ (*P* < 0.01) (**Figure 7C**), suggesting that *Hp* infection increases the proliferation of dysplastic glands.

**Figure 7.**
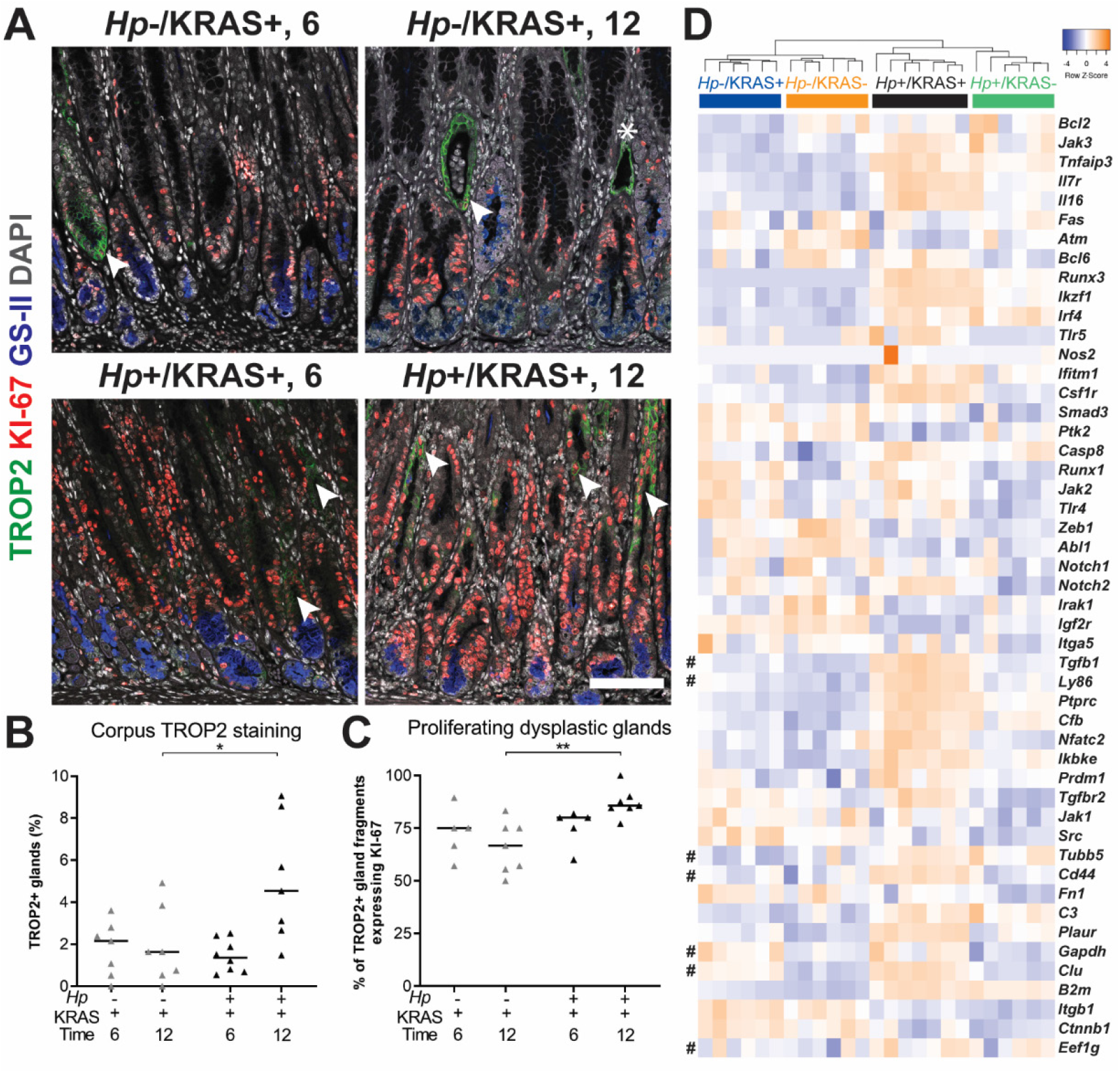
*Hp* infection increases dysplasia and cancer gene expression in KRAS+ mice. Stomachs from *Hp*-/KRAS+ and *Hp*+/KRAS+ mice were assessed through immunofluorescence microscopy (**A-C**) and gene expression analysis (**D**). (**A-C**) Corpus tissue from *Hp*-/KRAS+ and *Hp*+/KRAS+ mice obtained after six or 12 weeks (N=2 independent mouse experiments per time point and n=5-8 mice per group) was stained with antibodies against TROP2 (green) and KI-67 (red), the lectin GS-II (blue) and DAPI (grey). (**A**) Representative images are shown, arrows show TROP2+/KI-67+ gland fragments, and the asterisk shows a TROP2+/KI-67-gland fragment. Scale bar, 100 µm. (**B**) TROP2+ gland fragments were enumerated as a percentage of total gland fragments detected. (**C**) TROP2+ glands or gland fragments were assessed for KI-67 staining and the percentage of dual-positive gland fragments is shown. Statistically significant comparisons are indicated by: * *P* < 0.05, ** *P* < 0.01, Kruskal-Wallis test with Dunn’s correction. (**D**) The expression of gastric cancer-associated genes discovered through literature searching is shown. RNA was extracted from stomach sections from *Hp*+/-, KRAS+/- mice at 12 weeks and gene expression was detected with the NanoString nCounter Mouse Immunology Panel. The dendrogram was produced by hierarchical clustering of gene expression and colored bars denote different treatment groups. Data comes from N=1 NanoString experiment with n=6-7 mice per group from N=2 independent mouse experiments. # denotes genes expressed in *Mist1-Kras* organoids ^41^.

We mined our NanoString gene expression data (**Figure 4A**) and identified 49 genes implicated in the development of gastrointestinal cancers ^41-46^. Hierarchical clustering analysis showed that as with the overall panel, *Hp* infection status had the greatest impact on expression of this subset of genes at 12 weeks, and that active KRAS expression also impacted gene expression, though to a lesser extent than *Hp* infection status did (**Figure 7D**). However, some genes, such as *Jak2, Notch2* and *Runx1*, were upregulated in KRAS+ mice regardless of infection status. Finally, metaplastic and dysplastic organoids generated from *Hp*-/KRAS+ mice at 12 and 16 weeks after active KRAS induction, respectively, were previously found to have unique phenotypes and gene expression signatures ^41^. Seven of these genes were found in our panel (**Figure 7D**, denoted with #). Expression of the metaplasia-associated gene *Clu* was strongly upregulated in KRAS+ mice, but the metaplasia-associated gene *Ly86* was only strongly expressed in *Hp*+/KRAS+ mice. Of the dysplasia-associated genes, *Tubb5* was elevated in *Hp*+ mice, *Gapdh* was elevated in KRAS+ mice, and *Eef1g* was not differentially expressed among treatment groups. Finally, *Cd44* and *Tgfb1* were previously found in both metaplastic and dysplastic organoids ^41^ and in our study were strongly elevated in *Hp*+/KRAS+ mice at 12 weeks relative to the other mouse groups. Thus, gene expression in *Hp*+/KRAS+ whole stomachs at 12 weeks is distinct from *Hp*-/KRAS+ organoids generated at either 12 or 16 weeks, further supporting the hypothesis that concomitant *Hp* infection and active KRAS expression results in a unique gastric environment that is distinct from either *Hp*-/KRAS+ or *Hp*+/KRAS-mice.

### *Sustained Hp* infection is necessary to elicit changes to metaplasia, dysplasia and cell proliferation

Finally, we tested the impact of antibiotic eradication of *Hp* in two lines of experiments (**Figure 8 and Supplementary Figure S12**). First mice were infected with *Hp* or mock-infected, and active KRAS was induced. Starting at two weeks after active KRAS induction, mice received two weeks of “triple therapy” of tetracycline, metronidazole and bismuth ^47^ or vehicle (water) as a control, and were euthanized at six weeks (**Figure 8A and B**). Notably, *Hp*+/KRAS+ mice that received triple therapy had low CD44v (**Figure 8A and C**) and high TFF3 (**Figure 8A and D**) expression, similar to *Hp*-/KRAS+ mice (whether untreated as in **Figure 6E-F**, or treated with triple therapy or water). Thus, eradication of *Hp* early in the course of infection prevents the altered course of metaplasia seen at six weeks. Similarly, we tested the effects of triple therapy after six weeks (a time point where significant *Hp*-dependent changes to the tissue are already evident) with euthanasia at 12 weeks (**Figure 8E and F**). *Hp*+/KRAS+ mice that received triple therapy had reduced TROP2+ glands at 12 weeks (**Figure 8E and G**), suggesting that sustained *Hp* presence is necessary to accelerate dysplasia in this model. Finally, we found that at both time points antibiotic-treated *Hp*+/KRAS+ mice had reduced KI-67 staining (**Figure 8B, F and H**), demonstrating that sustained *Hp* presence is also required for the hyperproliferation phenotype in *Hp*+/KRAS+ mice.

**Figure 8.**
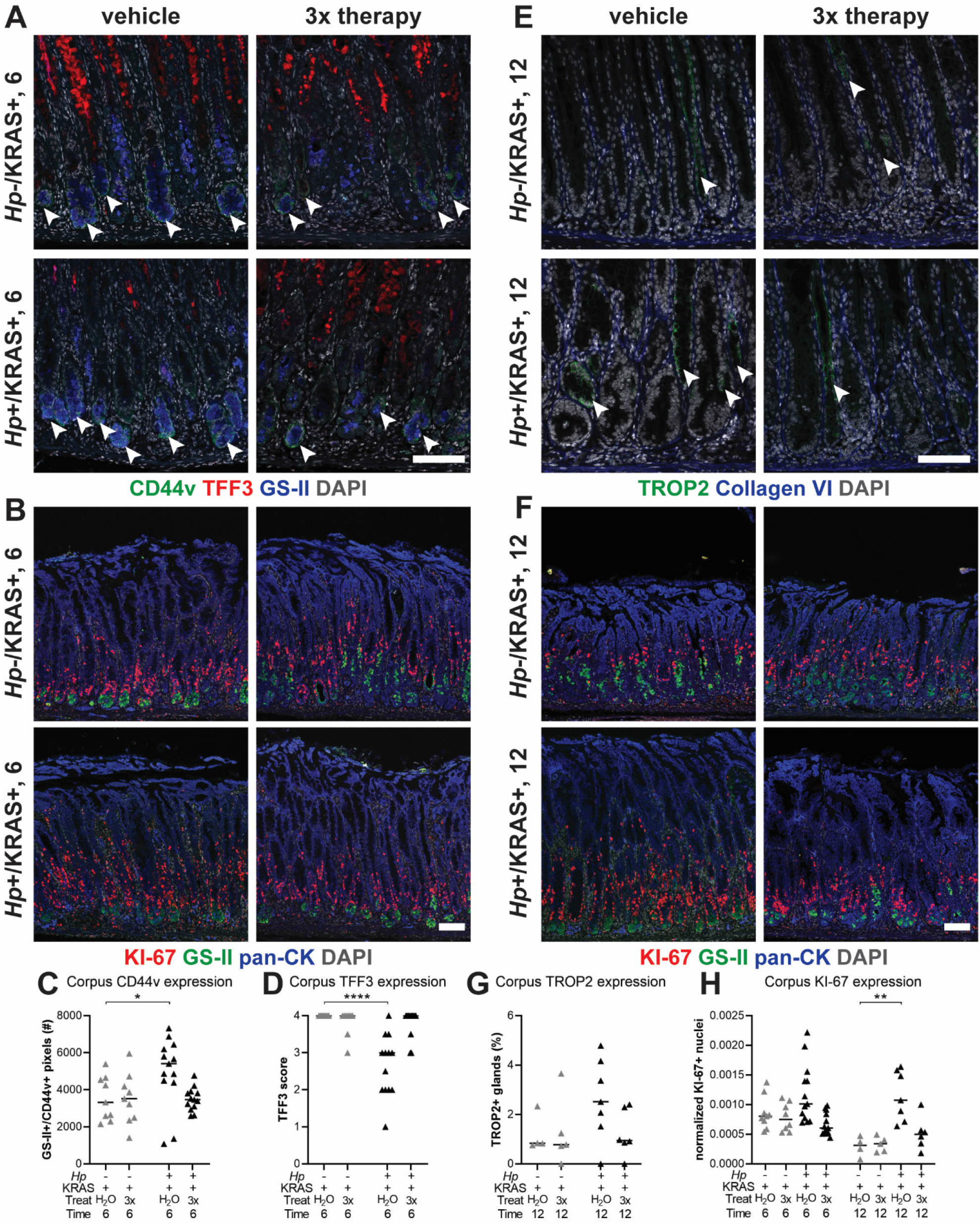
Antibiotic eradication prevents *Hp*-associated changes in *Hp*+/KRAS+ mice. First, mice were infected with *Hp* or mock-infected, then active KRAS was induced. Starting at two weeks (**A-D**) or six weeks (**E-G**), mice received two weeks of daily antibiotic therapy (triple therapy of tetracycline, metronidazole and bismuth, “3x”) or vehicle control (“H_2_O”) Mice were euthanized after six (**A-D**) or 12 (**E-G**) weeks and immunofluorescence microscopy was used to assess tissue phenotypes. (**A**) Stomachs were stained with antibodies against CD44v (green, arrowheads) and TFF3 (red), the lectin GS-II (blue) and DAPI (grey) in N=1 staining experiment. (**E**) Stomachs were stained with antibodies against TROP2 (green, arrowheads) and collagen VI (blue) and DAPI (grey) in N=1 staining experiment. (**B and F**) Stomachs were stained with antibodies against KI-67 (red) and pan-cytokeratin (blue), the lectin GS-II (green) and DAPI (grey) in N=2 staining experiments. (**C, D, G, H**) Three to five representative images per mouse were quantitatively or semi-quantitatively assessed and the median value for each mouse is plotted. Bars on the graphs indicate the median value for each mouse group. (**C**) The number of GS-II+/CD44v+ pixels per image was quantified. (**D**) TFF3 staining was semi-quantitatively scored in a blinded fashion. Non-specific staining was not included in the score (see **Figure S9**). (**G**) The percentage of Trop+ gland fragments was determined by collagen VI staining to detect individual gland fragments. (**H**) KI-67+/DAPI+ nuclei were enumerated and normalized to the DAPI content (total number of DAPI+ pixels) of each image. Statistically significant comparisons are indicated by: * *P* < 0.05, ** *P* < 0.01, **** *P* < 0.0001, Kruskal-Wallis test with Dunn’s correction. Representative images are shown. Scale bars, 100 µm.

## Discussion

In this study we examined the effect of *Hp* infection in stomachs expressing active KRAS (**Table 1**). Up to 40% of human gastric cancers have genetic signatures of RAS activity ^24, 25^. Activation of RAS and/or other oncogenic pathways in humans could be a consequence of *Hp*-driven inflammation. In our model, KRAS activation serves as a tool to model the consequences of oncogenic inflammation caused by *Hp* infection. Because active KRAS alone is sufficient to cause gastric preneoplastic progression ^21^, it might be expected that *Hp* infection would have no impact on KRAS-driven phenotypes. However, we found that *Hp* infection did influence preneoplastic progression in this model. *Hp* infection in KRAS-expressing mice led to more severe inflammation, an altered trajectory of metaplasia, substantial cell proliferation, and increased dysplasia compared to active KRAS alone. Additionally, eradication of *Hp* with antibiotics prevented these tissue changes, in accordance with a major long-term study of *Hp* eradication in Colombian adults with precancerous lesions, which showed that continuous *Hp* presence was significantly associated with disease progression ^48^. Thus, our study supports the hypothesis that sustained *Hp* can impact the molecular course of cancer development, beyond just initiating chronic inflammation.

**Table 1.** Summary of differences between *Hp*-/KRAS+ and *Hp*+/KRAS+ mice. HAI, histological activity index.

| Parameter | Hp-/KRAS+ | Hp+/KRAS+ | Relevance to Humans |
| --- | --- | --- | --- |
| Tissue histology | Inflammation, loss of parietal cells, surface epithelial cell hyperplasia, tortuous glands. Median HAI is 13 at six weeks and 11 at 12 weeks | Worsening of these parameters. Median HAI is 19 at six weeks and 20.5 at 12 weeks | Inflammation, parietal cell loss and metaplasia are hallmarks of gastric preneoplastic progression <sup>4, 5</sup> |
| Immune gene expression | Less upregulation of T cell and macrophage-related genes | Upregulation of genes related to T cells and T cell exhaustion, macrophages and gastric cancer; unique inflammatory gene signature | Many genes upregulated in *Hp*+/KRAS+ mice were shown to be upregulated in human gastrointestinal cancers <sup>41-46</sup> |
| T cells | Moderate T cells, predominantly CD4+. T cells mostly restricted to the base of the glands | Many T cells, predominantly CD4+, including FOXP3+. T cells extend throughout the glands | *Hp* infection leads to recruitment and activation of T cells <sup>14-16</sup>, which may lead to mucosal damage and accumulation of mutations <sup>17, 18</sup><br>Exhausted T cells may be targeted by immune checkpoint inhibitors <sup>58</sup> |
| Macrophages | F4/80+/CD163+ (M2) cells are present, with a few F4/80+/MHC class II+ (M1) as well | Some F4/80+/MHC class II+ (M1) cells are present, with very few F4/80+/CD163+ (M2) | M2 macrophages are found in human SPEM and IM <sup>34</sup><br>Macrophages can infiltrate into gastric tumors and are associated with worse surgical outcomes <sup>62</sup> |
| Metaplasia marker expression (6 weeks) | Moderate CD44v9, high TFF3, low MUC2 | High CD44v9, moderate TFF3, low MUC2 | SPEM (CD44v10+) and IM (MUC2+, TFF3+) are both precursor lesions associated |
| Metaplasia marker expression (12 weeks) | Moderate CD44v9, high TFF3, low MUC2 | High CD44v9, low TFF3, low MUC2 | with human gastric adenocarcinoma <sup>50-52</sup> |
| Cell proliferation | Moderate numbers of KI-67+ cells near the base of the glands | Far more of KI-67+ cells, near the base and throughout the glands | Cell proliferation was greater in IM than in gastritis or healthy human stomachs <sup>37</sup> |
| Dysplasia | Low TROP2 expression; most TROP2+ glands are proliferative (KI-67+) | Moderate TROP2 expression at 12 weeks and almost all TROP2+ glands are proliferative (KI-67+) | TROP2 overexpression is associated with many cancers <sup>38, 39</sup> and was recently shown to be a gastric dysplasia marker in humans <sup>40</sup> |
| Antibiotic therapy | No impact on metaplasia, dysplasia or cell proliferation marker expression | *Hp* eradication prevents altered metaplasia, accelerated dysplasia and cell proliferation phenotypes; resembles *Hp*-/*KRAS*+ | *Hp* eradication prevented disease progression in patients with precancerous lesions <sup>48</sup><br>In cancer patients with *Hp*, eradication reduces metachronous cancer <sup>12</sup> |

Different mouse models exhibit different clinical features of preneoplastic progression: for example, *Helicobacter* infection alone causes SPEM, but does not cause foveolar hyperplasia or IM in C57BL/6 mice, whereas uninfected *Mist1-Kras* mice exhibit all of these tissue states after active KRAS induction ^49^. Here we found that the combination of sustained *Hp* infection and active KRAS expression has a unique impact on the development of gastric metaplasia that is not observed with either individual parameter. Compared to *Hp*-/KRAS+ mice, *Hp*+/KRAS+ mice had no difference in foveolar hyperplasia or expression of the IM marker MUC2, but had decreased expression of the IM marker TFF3 and increased expression of the SPEM marker CD44v. Further work is needed to determine whether these changes in metaplasia marker expression may reflect increased SPEM vs. a process similar to incomplete IM. To our knowledge, of the various mouse models of gastric corpus preneoplastic progression, only *Mist1-Kras* mice exhibit true IM (indicated by TFF3+ and MUC2+ glands) in 100% of mice; most other models exhibit SPEM with or without intestinalizing characteristics ^49^. The finding that *Hp* infection altered TFF3 expression in *Mist1-Kras* mice is therefore quite striking. TFF3 expression was moderate in both treatment groups at two weeks, and it is not yet known whether TFF3 expression may have peaked in *Hp*+/KRAS+ mice at an intermediate time point, such as four weeks, or was never as strongly expressed as in *Hp*-/KRAS+ mice. Notably, several human studies have reported that the association of SPEM with gastric adenocarcinoma is equal to or even greater than that of IM ^50-52^, leading to questions in the field regarding the trajectory of metaplasia development prior to gastric cancer onset. Our findings suggest that the trajectory of metaplasia could differ depending on whether or not *Hp* remains present in the stomach throughout preneoplastic progression.

Sustained *Hp* infection also caused a striking increase in cell proliferation as indicated by KI-67 staining, and increased TROP2 staining at 12 weeks. TROP2 expression was lower in our mice than what was previously reported ^40^, which may reflect differences in animal housing conditions, different methods of tissue fixation and processing, or components of the microbiome (although antibiotic perturbation in *Hp*-/KRAS+ mice had no effect on expression of TROP2 or metaplasia markers). Nonetheless, within our controlled experiments, TROP2 staining was greatest in *Hp*+/KRAS+ mice, suggesting that infection accelerates the onset of dysplasia. While almost all of the TROP2+ glands in *Hp*+/KRAS+ mice had co-localized KI-67 staining, demonstrating proliferation of dysplastic glands, there were also many KI-67+ cells in TROP2-glands. Future studies will seek to elucidate the specific cell types that are proliferating in *Hp*+/KRAS+ mice. Despite the enhanced proliferation of dysplastic glands, *Hp*+/KRAS+ mice did not develop gastric tumors within 12 weeks. One limitation of our study is that the Mist1 promoter is expressed outside the stomach in secretory lineages, including the salivary glands. Approximately four months after active KRAS induction, *Mist1-Kras* mice require humane euthanasia due to salivary gland tumors. Thus, we cannot test whether *Hp* infection promotes even more severe phenotypes, such as tumor development, at later time points. Specifically targeting active KRAS to the chief cells via other promoters could overcome this hurdle. However, *Hp*+/KRAS+ mice did have increased expression of genes known to be associated with gastrointestinal cancers. It may be that at least some of these genes are associated with gastric cancer simply because they reflect *Hp* infection, the biggest risk factor for gastric cancer development.

Interestingly, we found that a few *Hp*+/KRAS+ mice naturally cleared their infection, yet still had a high degree of immunopathology. A limitation of modeling *Hp* infection in mice is the inability to monitor bacterial burdens over time. It may be that the mice in question cleared their infection just prior to euthanasia, with no time for *Hp*-driven tissue changes to reverse. Alternatively, *Hp* infection may lead to a “point of no return,” after which immunopathology develops even in the absence of *Hp*. This hypothesis has been used to explain the lack of detectable *Hp* in about half of human gastric tumors ^53, 54^. However, we found that antibiotic eradication of *Hp* after six weeks prevented the accelerated dysplasia and hyperproliferation phenotypes at 12 weeks. Thus, if a “point of no return” exists in this model, it must occur after six weeks.

We note that no significant differences in metaplasia or dysplasia marker expression were observed between *Hp*-/KRAS+ mice and *Hp*+/KRAS+ mice at two weeks, suggesting that the adaptive immune response to *Hp* infection may be what promotes the differences in metaplasia and dysplasia observed at later time points. This observation agrees with previous findings that T cells were necessary for *Helicobacter*-associated gastritis ^27, 28^ and metaplasia development ^55^. The immune response seen in *Hp*+/KRAS+ mice at six and especially 12 weeks far exceeded what was observed in either *Hp*+/KRAS- or *Hp*-/KRAS+ mice, and indeed is much greater than what is typically seen in *Hp* mouse models. Notably, this inflammation did not eradicate *Hp*: most *Hp*+/KRAS+ mice remained colonized at 12 weeks. *Hp* cells were observed within the lumen of metaplastic glands, where they may be protected from direct immune cell interaction. Additionally, *Hp* has multiple strategies to prevent immune-mediated clearance ^13^, including disruption of normal T cell functions by: triggering upregulation of PD-L1, a T cell inhibitory ligand that binds programmed cell death protein-1 (PD-1), on gastric epithelial cells, leading to T cell exhaustion ^19^; inducing anergy through promoting T cell expression of the CTLA-4 co-receptor ^56^; inhibiting T cell proliferation and normal effector functions with the vacuolating cytotoxin VacA ^20^; and the induction of tolerogenic dendritic cells, which promote the differentiation of naive T cells into immunosuppressive regulatory T cells ^13^. It remains unknown whether and to what extent *Hp* may disrupt T cell functions in our model. *Hp*+/KRAS+ mice were unique in their upregulation of *Foxp3* and had FOXP3+ T cells at 12 weeks, but these cells were not sufficient to limit immunopathology. As well, in *Hp*+/KRAS+ mice we observed strong transcriptional upregulation of T cell exhaustion-related genes, such as *Pdcd1* (PD-1), and *Ctla4*, implicated in T cell anergy. By immunohistochemistry we saw evidence of PD-1 expression in both Hp-/KRAS+ and Hp+/KRAS+ mice. Further studies are needed to characterize the exact nature of immune cell polarization differences between treatment groups; to confirm whether the T cells observed in *Hp*+/KRAS+ mice may be exhausted, anergic or senescent; and to test whether immunosuppression or immunomodulation would be protective against *Hp*’s effects in the model. Nonetheless, it is clear that the combination of *Hp* infection and active KRAS expression leads to a potent and unique inflammatory state. Given that immunotherapy is still under-utilized in gastric cancer ^57^ and only a subset of patients benefit from such treatments ^58^, a better understanding of how active *Hp* infection may alter the immune microenvironment during gastric metaplasia and cancer development is urgently needed and may lead to new therapeutic strategies.

When *Hp* was first discovered to be a bacterial carcinogen, studies using tissue histology rarely detected *Hp* within gastric tumors. Such studies may have helped establish the belief that although *Hp* initiates the gastric cancer cascade, by the time gastric cancer is developed, *Hp* no longer matters – the so-called “hit-and-run” model. However, more sensitive methods detect *Hp* in about half of gastric tumors ^9-11^, indicating that a large percentage of patients maintain active *Hp* infection throughout cancer development. Notably, *Hp* eradication combined with endoscopic resection of early gastric cancer significantly prevents metachronous gastric cancer ^12^. As well, a recent study of 135 *Hp*-seropositive subjects with IM found that patients with active *Hp* infection (determined by histology and/or sequencing) were significantly more likely to have somatic copy number alterations (sCNA), and that patients with sCNA were more likely to experience IM progression ^42^. Given these observations, there is an urgent need for preclinical models that identify unique features of gastric neoplasia with vs. without concomitant *Hp* infection, both to understand the etiology of gastric cancer and to determine the impact of infection on different therapeutic approaches. We have shown here that *Hp* can significantly impact metaplasia and dysplasia development in a clinically relevant mouse model, which suggests that gastric preneoplastic progression can develop differently in the presence vs. absence of *Hp*. Future studies will elucidate the molecular mechanism(s) through which *Hp* exerts its effects in this model, and test whether active *Hp* infection during metaplasia or cancer may represent a therapeutic vulnerability that could be targeted with immunotherapy.

## Materials and Methods

### Ethics Statement

All mouse experiments were performed in accordance with the recommendations in the National Institutes of Health Guide for the Care and Use of Laboratory Animals and were approved by the Fred Hutch Institutional Animal Care and Use Committee, protocol number 1531.

### Helicobacter pylori *Strains and Growth Conditions*

*Hp* strain PMSS1, which is also called 10700 and which is CagA+ with an active type IV secretion system ^47, 59^, and derivatives were cultured at 37°C with 10% CO_2_ and 10% O_2_ in a trigas incubator (MCO-19M, Sanyo). *Hp* titers were determined by quantitative culture on horse blood Columbia agar (BD Biosciences) with antibiotic supplementation to prevent overgrowth of commensal organisms ^60^.

### Mist1-Kras *Mouse Model*

Mist1-CreERT2 Tg/+; LSL-K-Ras (G12D) Tg/+ (“*Mist1-Kras*,” C57BL/6 background) mice were described previously ^21^. Eight to 16 week-old male and female mice were infected with 5×10^7^ mid-log culture *Hp* cells in 100 µL of liquid media (BB10) containing 90% (v/v) Brucella broth (BD Biosciences) and 10% fetal bovine serum (Gibco), or mock-infected with 100 µL of BB10. To induce active (oncogenic) KRAS expression, mice received three subcutaneous doses of 5 mg of tamoxifen (Sigma) in corn oil (Sigma) over three days, or were sham-induced with corn oil, starting one day after *Hp* or mock infection. We used n=10-16 mice per group in N=2 independent experiments per time point, except for antibiotic eradication after six weeks, which used n=5-7 mice per group in N=1 experiment. Subsequent analyses of tissue changes used n=5-12 samples per treatment group, chosen randomly. Mice were humanely euthanized by CO_2_ inhalation followed by cervical dislocation at two, six or 12 weeks after infection and transgene induction. Stomachs were aseptically harvested; a portion was used for *Hp* culture and the rest fixed in 10% neutral-buffered formalin for sectioning. For antibiotic eradication, mice received 4.5 mg/mL metronidazole, 10 mg/mL tetracycline hydrochloride and 1.2 mg/mL bismuth subcitrate, or vehicle (water), by oral gavage for two weeks ^47^.

### Histology

A veterinary pathologist (A.K.) scored hematoxylin and eosin-stained tissue sections in a blinded fashion according to criteria adapted from Rogers ^26^. The sum of the individual scores for each criterion were summed to generate a histological activity index (HAI) score.

### Gene Expression Analysis

RNA was extracted from five 4-µm formalin-fixed, paraffin-embedded (FFPE) stomach sections per mouse using the AllPrep DNA/RNA FFPE Kit (Qiagen) and gene expression was detected using the nCounter Mouse Immunology Panel (NanoString). Gene expression differences were detected using nSolver software (NanoString) (**Supplementary Table S2**). Hierarchical clustering was performed and heat maps were generated with HeatMapper ^61^ using the average linkage method with Euclidian distance, using log_2_-transformed gene expression data.

### Multiplex Immune Immunohistochemistry

Tissues were stained using the Leica BOND RX system using Leica BOND reagents for dewaxing, antigen retrieval/antibody stripping, and rinsing after each step (**Supplementary Table S1**). To obtain multiplex labeling, one primary antibody was applied for 60 minutes, followed by the relevant secondary antibody and OPAL fluorophore for 10 minutes each. Slides were stripped of excess antibodies and the next primary and secondary were applied in sequence until all eight primary antibodies were applied.

### Immunofluorescence Microscopy and Quantitation of Staining

Immunofluorescence microscopy to assess gastric preneoplastic progression was performed as previously described ^21^ (**Supplementary Table S3**). Three to five representative images of corpus tissue per mouse were used for quantitation and the median value was reported for each mouse. Scripts for quantitation of KI-67, GS-II, CD44v and TROP2 can be found on GitHub at: https://github.com/salama-lab/stomach-image-quantitation.

### Statistical Analyses

For NanoString analysis, *P*_adjusted_ values were generated in nSolver with the Benjamini-Yekutieli procedure for controlling the false discovery rate. Other statistics were performed in GraphPad Prism v7.01. Comparisons of three or more groups were performed with the Kruskal-Wallis test with Dunn’s multiple test correction. *P* < 0.05 was considered statistically significant.

## Acknowledgement

The authors wish to acknowledge Savanh Chanthaphavong and Louis Kao for assistance with experimental histopathology and Alicia M. Meyer for assistance with mouse experiments.

## Supplementary Material

Expanded Materials and Methods

Supplementary Figures 1-7

Supplementary Tables 1 and 3

Supplemental References

## Expanded Materials and Methods

### Ethics Statement

All mouse experiments were performed in accordance with the recommendations in the National Institutes of Health Guide for the Care and Use of Laboratory Animals. The Fred Hutchinson Cancer Research Center is fully accredited by the Association for Assessment and Accreditation of Laboratory Animal Care and complies with the United States Department of Agriculture, Public Health Service, Washington State, and local area animal welfare regulations. Experiments were approved by the Fred Hutch Institutional Animal Care and Use Committee, protocol number 1531.

### Helicobacter pylori *Strains and Growth Conditions*

*Helicobacter pylori* strain PMSS1, which is also called 10700 and which is CagA+ with an active type IV secretion system ^1, 2^, and derivatives were cultured at 37°C with 10% CO_2_ and 10% O_2_ in a trigas incubator (MCO-19M, Sanyo). Cells were grown on solid media containing 4% Columbia agar (BD Biosciences), 5% defibrinated horse blood (HemoStat Laboratories) 0.2% β-cyclodextrin (Acros Organics), 10 μg/ml vancomycin (Sigma), 5 μg/ml cefsulodin (Sigma), 2.5 U/mL polymyxin B (Sigma), 5 μg/ ml trimethoprim (Sigma) and 8 μg/ml amphotericin B (Sigma). For mouse infections, bacteria grown on horse blood plates were used to inoculate liquid media (BB10) containing 90% (v/v) Brucella broth (BD Biosciences) and 10% fetal bovine serum (Gibco), which was cultured shaking at 200 rpm overnight and grown to an optical density at 600 nm of 0.4 - 0.6 (mid-log phase), from which an inoculum of approximately 5×10^7^ *Hp* cells per 100 µl BB10 was prepared. To determine *Hp* titers in the stomach, harvested tissues were weighed, serially diluted, and plated on the solid media described above, with the addition of bacitracin (200 μg/ml, Acros Organics) to prevent growth of the stomach microbiota.

### Mist1-Kras *Mouse Model*

A breeding pair of Mist1-CreERT2 Tg/+; LSL-K-Ras (G12D) Tg/+ (“*Mist1-Kras*”) mice on the C57BL/6 background, described previously ^3^, was obtained from Vanderbilt University (E.C. and J.R.G) and used to establish a colony at Fred Hutchinson Cancer Research Center. Mice were housed two to five per cage, with cages docked in HEPA-filtered ventilation racks that provide airflow control on a 12 hour light/dark cycle, and had access to chow (LabDiet) and water ad libitum. At weaning, ear punches were collected and used for genotyping as previously described ^3^ with the following primers: Mist1 WT F: CCAAGATCGAGACCCTCACG; Mist1 WT R: ACACACACAGCCCTTAGCTC Mist1 Cre F: ACCGTCAGTACGTGAGATATCTT; Mist1 Cre R: CCTGAAGATGTTCGCGATTATCT; active KRAS F: TCTCTGCAGTTGTTGGCTCCAAC; active KRAS R: GCCTGAAGAACGAGATCAGCAGCC. Healthy eight to 16 week-old male and female mice (randomly allocated to treatment groups) were infected with 5 × 10^7^ mid-log culture *Hp* cells in 100 µL of BB10, or mock-infected with 100 µL of BB10, via oral gavage. To induce active (oncogenic) KRAS expression, mice received three subcutaneous doses of 5 mg of tamoxifen (Sigma) in corn oil (Sigma) over three days, or were sham-induced with corn oil, starting one day after *Hp* or mock infection. For antibiotic eradication, mice received “triple therapy” of 4.5 mg/mL metronidazole, 10 mg/mL tetracycline hydrochloride and 1.2 mg/mL bismuth subcitrate, or vehicle (water), by oral gavage.^1^ Mice received six doses in seven days, two weeks in a row. Mice were humanely euthanized by CO_2_ inhalation followed by cervical dislocation two, six or 12 weeks after infection and transgene induction. Stomachs were aseptically harvested and most of the forestomach (non-glandular region) was discarded, leaving only the squamocolumnar junction between the forestomach and glandular epithelium. Approximately one-third of the stomach was homogenized and plated to enumerate *Hp*. The remaining approximately two-thirds of the stomach was fixed in 10% neutral-buffered formalin phosphate (Fisher), then embedded in paraffin and cut into 4 µm sections on positively-charged slides. *Hp*+/KRAS+ and *Hp*-/KRAS mice did not exhibit overall health differences; body weights and behaviors were similar at time of euthanasia.

### Histology

Stomach sections were stained with hematoxylin and eosin (H&E). A veterinary pathologist (A.K.) scored the slides in a blinded fashion according to criteria adapted from Rogers ^4^. Corpus tissue was evaluated for inflammation, epithelial defects, oxyntic atrophy, hyperplasia (tissue thickness), hyalinosis, pseudopyloric metaplasia, mucous metaplasia and dysplasia. The sum of the individual scores for each criterion were summed to generate a histological activity index (HAI) score. Scoring criteria are described below. HAI was not correlated with sex or with age of mice at sacrifice.

#### Inflammation

Multifocal aggregates of inflammatory cells merit a score of 1. As the aggregates coalesce across multiple high-power fields (40X objective), the score increases to 2. Sheets of inflammatory cells and/or lymphoid follicles in the mucosa or submucosa receive a score of 3. Florid inflammation that extends morally or transmurally is a score of 4.

#### Epithelial defects

A tattered epithelium with occasional dilated glands is a score of 1. As the epithelium becomes attenuated and ectatic glands become more numerous, the score increases to 2. Inapparent epithelial lining of the surface with few recognizable gastric pits are given a score of 3. Score 4 is reserved for mucosal erosions.

#### Oxyntic atrophy

The oxyntic mucosa is defined by the presence of chief and parietal cells. Loss of up to half of the chief cells merit a score of 1. In instances with near complete loss of chief cells and minimal loss of parietal cells, a score 2 is assigned. The absence of chief cells with half the expected number of parietal cells is given a score of 3. Score 4 signifies near total loss of both chief and parietal cells.

#### Surface epithelial hyperplasia

This score indicates elongation of the gastric gland due to increased numbers of surface (foveolar) and/or antral-type epithelial cells. Relative to the expected length of a normal gastric pit, a score of 1 indicates a 50% increase in length. A score of 2 is twice the expected length, a score of 3 is three times the expected length, and a score of 4 is four times the expected length.

#### Hyalinosis

This mouse-specific gastritis lesion refers to the presence of brightly eosinophilic round or crystalline structures in the murine gastric surface epithelium. The presence of epithelial hyalinosis is given a score of 1 while absence of hyalinosis is a score 0.

#### Pseudopyloric metaplasia

Pseudopyloric metaplasia is the loss of oxyntic mucosa and replacement with glands of a more antral phenotype. The score indicates the amount of replacement by antralized glands. Less than 25% replacement is a score of 1, 26-50% replacement is a score of 2, 51-75% replacement is a score of 3, and greater than 75% replacement is a score of 4.

#### Mucous metaplasia

This mouse-specific gastritis lesion is defined as the replacement of oxyntic cells with mucous producing cells that resemble Brunner’s glands of the duodenum. The score is assigned based on the percentage of mucosa affected. A score 1 indicates less than 25% involvement, a score of 2 indicates 26-50% involvement, a score of 3 is 51-75% involvement, and a score of 4 means that greater than 75% of the mucosa is involved.

#### Dysplasia

Dysplasia indicates a cellular abnormality of differentiation. In score 1 lesions, the glands are elongated with altered shapes, back-to-back forms, and asymmetrical cellular piling. In score 2, the dysplastic glands may coalesce with glandular ectasia, branching, infolding, and piling of cells. Gastric intraepithelial neoplasia (GIN) is given a score of 3 and invasive carcinoma is a score of 4. The dysplasia score describes the most severe lesion(s) in each mouse.

### Gene Expression Analysis

RNA was extracted from five 4-µm FFPE stomach sections per mouse using the AllPrep DNA/RNA FFPE Kit (Qiagen) and gene expression was detected using the nCounter Mouse Immunology Panel (NanoString). Gene expression differences were detected using nSolver software (NanoString) and are given in **Supplementary Table 2**. Volcano plots were constructed by taking the log_2_ of the fold change and the -log_10_ of the unadjusted *P* value for each gene. The *P*_adjusted_ lines show genes meeting the threshold for significance after correction with the Benjamini-Yekutieli procedure for controlling the false discovery rate. Hierarchical clustering was performed and heat maps were generated through HeatMapper ^5^ using the average linkage method with Euclidian distance, with log_2_-transformed gene expression data. Clustering was applied to rows (genes) and columns (mice). To identify the unique gene signature in *Hp*+/KRAS+ mice, gene expression values were normalized to the geometric mean of the expression in *Hp*-/KRAS-mice, and all genes were identified for which the geometric mean of the fold change in *Hp*+/KRAS+ mice was >2 and in *Hp*+/KRAS- and *Hp*-/KRAS+ mice was <1.5, or the geometric mean of the fold change in *Hp*+/KRAS+ mice was <0.5 and in *Hp*+/KRAS- and *Hp*-/KRAS+ mice was >0.667.

### Multiplex Immunohistochemistry for Immune Cell Detection

Slides were baked for 60 minutes at 60°C and then dewaxed and stained on a Leica BOND RX system (Leica, Buffalo Grove, IL) using Leica BOND reagents for dewaxing (Dewax Solution), antigen retrieval/antibody stripping (Epitope Retrieval Solution 2), and rinsing (Bond Wash Solution). Antigen retrieval and antibody stripping steps were performed at 100°C with all other steps at ambient temperature. Endogenous peroxidase was blocked with 3% H_2_O_2_ for 5 minutes followed by protein blocking with 10% normal mouse immune serum diluted in TCT buffer (0.05M Tris, 0.15M NaCl, 0.25% Casein, 0.1% Tween 20, 0.05% ProClin300 pH 7.6) for 10 minutes. Primary and secondary antibodies are given in **Supplementary Table 1**. The first primary antibody (position 1) was applied for 60 minutes followed by the secondary antibody application for 10 minutes and the application of the tertiary TSA-amplification reagent (PerkinElmer OPAL fluor) for 10 minutes. A high stringency wash was performed after the secondary and tertiary applications using high-salt TBST solution (0.05M Tris, 0.3M NaCl, and 0.1% Tween-20, pH 7.2-7.6). Undiluted, species-specific Polymer HRP was used for all secondary applications, either Leica’s PowerVision Poly-HRP anti-Rabbit Detection or ImmPress Goat anti-Rat IgG Polymer Detection Kit (Vector Labs) as indicated in Table S1. The primary and secondary antibodies were stripped with retrieval solution for 20 minutes before repeating the process with the second primary antibody (position 2) starting with a new application of 3% H_2_O_2_. The process was repeated until seven positions were completed. For the eighth position, following the secondary antibody application, Opal TSA-DIG was applied for 10 minutes, followed by the 20 minute stripping step in retrieval solution and application of Opal 780 fluor for 10 minutes with high stringency washes performed after the secondary, TSA DIG, and Opal 780 fluor applications. The stripping step was not performed after the final position. Slides were removed from the stainer and stained with DAPI for 5 minutes, rinsed for 5 minutes, and coverslipped with Prolong Gold Antifade reagent (Invitrogen/Life Technologies). Slides were cured overnight at room temperature, then whole slide images were acquired on the Vectra Polaris Quantitative Pathology Imaging System (Akoya Biosciences). The entire tissue was selected for imaging using Phenochart and multispectral image tiles were acquired using the Polaris. Images were spectrally unmixed using Phenoptics inForm software and exported as multi-image TIF files, which were analyzed with HALO image analysis software (Indica Labs). DAPI was used to detect individual cells and then cells expressing each marker were automatically detected based on signal intensity, and reported as a percentage of DAPI-positive cells.

### Immunofluorescence Microscopy of Epithelial Phenotypes

Stomach sections were prepared as described above. To validate the antibodies used to detect intestinal metaplasia, the entire intestinal tract from duodenum to colon was removed from an untreated C57BL/6 mouse. The cecum was discarded and the unflushed intestinal tract was rolled as a “Swiss roll,” fixed in 10% neutral-buffered formalin, paraffin-embedded and sectioned. Tissue sections were deparaffinized with Histo-Clear solution (National Diagnostics) and rehydrated in decreasing concentrations of ethanol. Antigen retrieval was performed by boiling slides in 10 mM sodium citrate (Fisher) or Target Retrieval Solution (Agilent Dako) in a pressure cooker for 15 minutes. Slides were incubated with Protein Block, Serum-Free (Agilent Dako) for 90 minutes at room temperature. Primary antibodies (**Supplementary Table 3**) were diluted in Protein Block, Serum Free, or Antibody Diluent, Background Reducing (Agilent Dako), and applied to the slides overnight at 4°C. Secondary antibodies were diluted 1:500 in Protein Block, Serum Free and slides were incubated for one hour at room temperature protected from light. Slides were mounted in ProLong Gold antifade reagent with DAPI (Invitrogen) and allowed to cure for 24 hours at room temperature before imaging. Slides were imaged on a Zeiss LSM 780 laser-scanning confocal microscope using Zen software (Zeiss) and three to five representative images of the corpus were taken.

### Quantitation of Staining

Three to five representative images of corpus tissue per mouse used for staining analysis and the median value was reported for each mouse. Investigators were blinded to the treatment groups. KI-67, GS-II, CD44v10 (orthologous to human CD44v9 and referred to herein as “CD44v”) and TROP2 markers were quantified from fluorescently immunolabelled tissue sections by custom-made scripts developed in MATLAB 2019a. Scripts can be found on Github at https://github.com/salama-lab/stomach-image-quantitation.

After background subtraction and denoising in each channel, positive pixels for DAPI, GS-II, CD44v or TROP2 were identified by image binarization using the Otsu method and morphological filtering. When appropriate, individual glands were segmented using the cytokeratin signal, which is predominant in glandular structures, or using the complement image of the collagen VI signal, which is excluded from glandular structures. For GS-II quantification, the fractional area of cytokeratin staining positive for GS-II was recorded. For TROP2 quantification, the fractional area of each gland fragment identified by collagen VI labelling was recorded, and gland fragments with ≥10% TROP2-positive pixels were considered TROP2-positive. To identify GS-II and CD44v double-positive regions, the GS-II binary mask was first dilated by a few pixels, since GS-II is cytoplasmic and CD44v is membrane-bound. The resulting number of overlapping pixels per image was then recorded. To assess KI-67 staining, individual KI-67-positive nuclei were identified using a watershed algorithm after distance transformation of the binarized signal, and then normalized by dividing by the total number of DAPI-positive pixels in the image.

TFF3, MUC2, and dual-positive TROP2/KI-67 staining were manually assessed. To assess TFF3 staining, images were scored manually using a semi-quantitative scale with the following criteria: 0 = no staining, 1 = 1-25% of glands are positive, 2 = 26-50% of glands are positive, 3 = 51-75% of glands are positive, 4 = >75% of glands are positive for TFF3. Positive TFF3 signal manifests as moderately bright, cell-associated staining with goblet cell-like morphology. Overly bright staining without distinct goblet-like morphology, and/or staining within the gland lumen (not cell-associated), was observed near the top of the glands in *Hp*-/KRAS-(healthy control) mice and was considered false-positive staining. To assess MUC2 staining, glands were detected by cytokeratin staining, and MUC2+ and MUC2-glands were manually counted. To assess gland fragments dual positive for TROP2 and KI-67, regions of TROP2+ staining that contained KI-67+ nuclei were counted and expressed as a percentage of all TROP2+ glands.

### Type IV Secretion System Activity

*Hp* strain PMSS1 was recovered from infected mice after euthanasia by serial dilution plating, described above, and five or six individual colonies per mouse were expanded and frozen at -80°C. Colonies were then grown and used in co-culture experiments with AGS cells (from a human gastric adenocarcinoma cell line; ATCC CRL-1739) as previously described ^6^. The input strain of PMSS1 (freezer stock) served as a positive control and a PMSS1Δ*cagE* mutant ^6^ served as a negative control. Infections were performed in triplicate and supernatants were collected after 24 hours of co-culture. IL-8 was detected using a human IL-8 enzyme-linked immunosorbent assay (ELISA) kit from BioLegend.

### Statistical Analyses

Volcano plots and heat maps were constructed from data generated by nSolver software (NanoString). Other statistics were performed in GraphPad Prism v7.01. Comparisons of three or more groups were performed with the Kruskal-Wallis test followed by Dunn’s multiple test correction. *P* < 0.05 was considered statistically significant. For histopathological evaluation of stomach sections and quantitation of staining, experimenters were blinded to the treatment groups.

**Figure S1.**
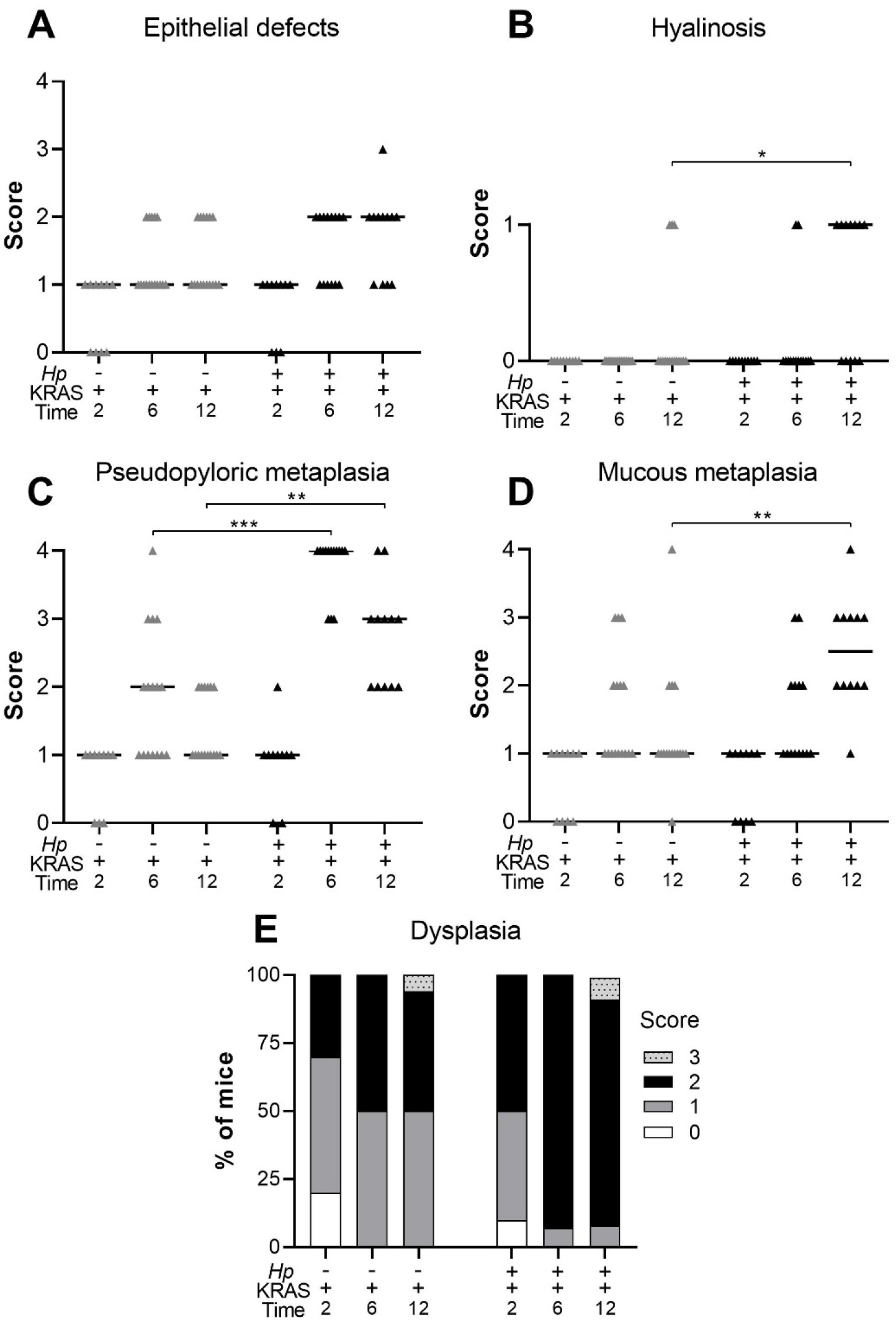
Concomitant *Hp* infection and active KRAS expression elicits changes to the stomach corpus. Corpus tissue from n=10-16 mice per group was evaluated for tissue pathology in a blinded fashion according to criteria adapted from Rogers ^4, 6^. Tissue was assessed for: (**A**) epithelial defects; (**B**) hyalinosis (a mouse-specific lesion denoted by the presence of brightly eosinophilic round or crystalline structures in the gastric surface epithelium); (**C**) pseudopyloric metaplasia (replacement of oxyntic mucosa with glands with a more antral-like phenotype); (**D**) mucous metaplasia (a mouse-specific lesion denoted by replacement of oxyntic mucosa with glands resembling Brunner’s glands of the duodenum); and (**E**) dysplasia, with gastric intraepithelial neoplasia given a score of 3. The sum of these subscores and those shown in the main text gives the histological activity index. Data are combined from N=2 independent mouse experiments per time point. Data points represent actual values for each individual mouse and bars indicate median values. Statistically significant comparisons are indicated by: * *P* < 0.05, ** *P* < 0.01, *** *P* < 0.001, Kruskal-Wallis test with Dunn’s multiple test correction.

**Figure S2.**
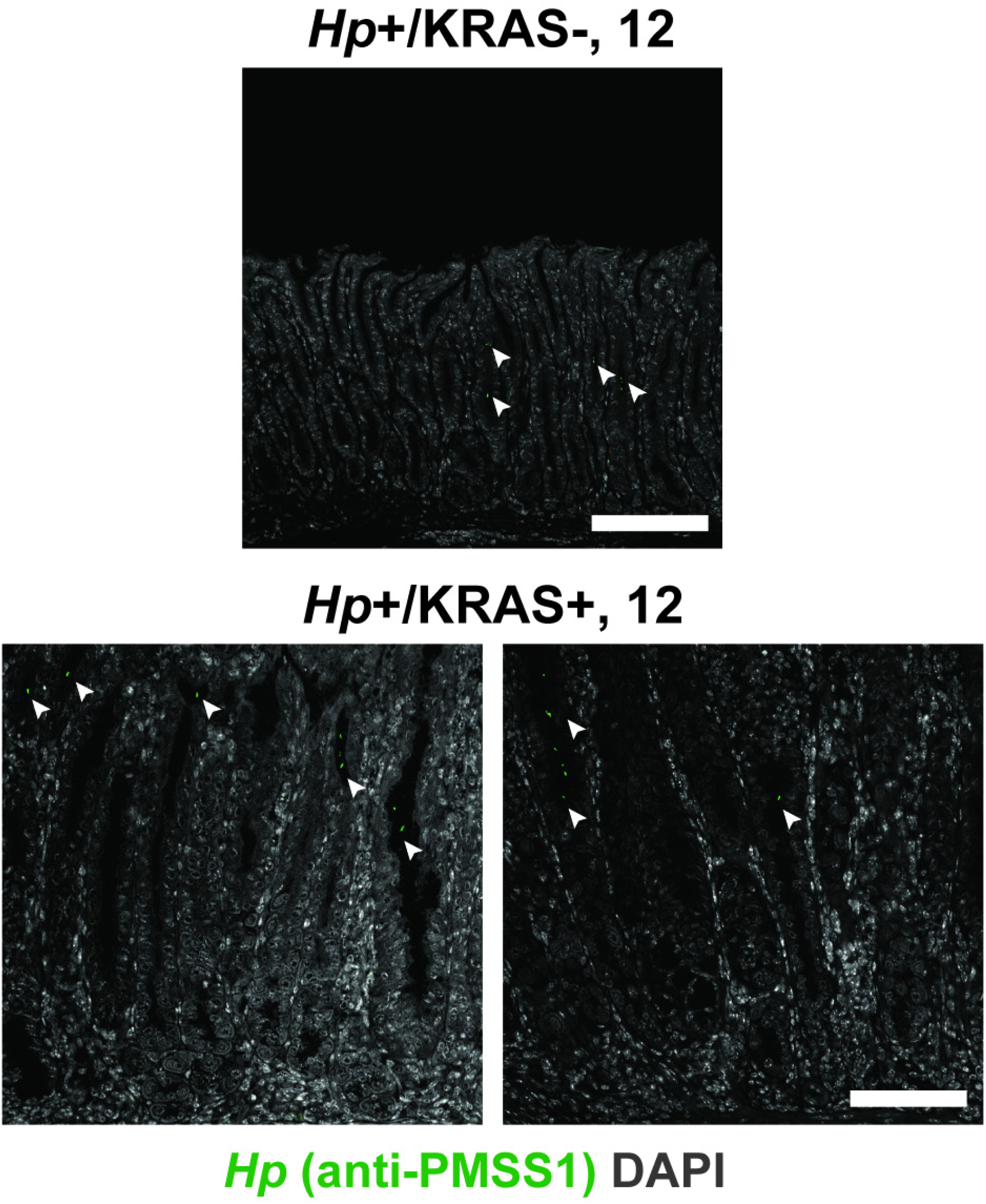
*Hp* is detected within gastric glands by immunohistochemistry. Thin stomach sections from *Hp*+/KRAS-mice (top) and *Hp*+/KRAS+ mice (bottom) obtained at the 12 week time point were stained with an anti-*Hp* strain PMSS1 antibody (green) and DAPI (grey) in N=1 staining experiment with n=4 mice per group. *Hp* (arrowheads) could be detected within the glands. Scale bars, 100 µm.

**Figure S3.**
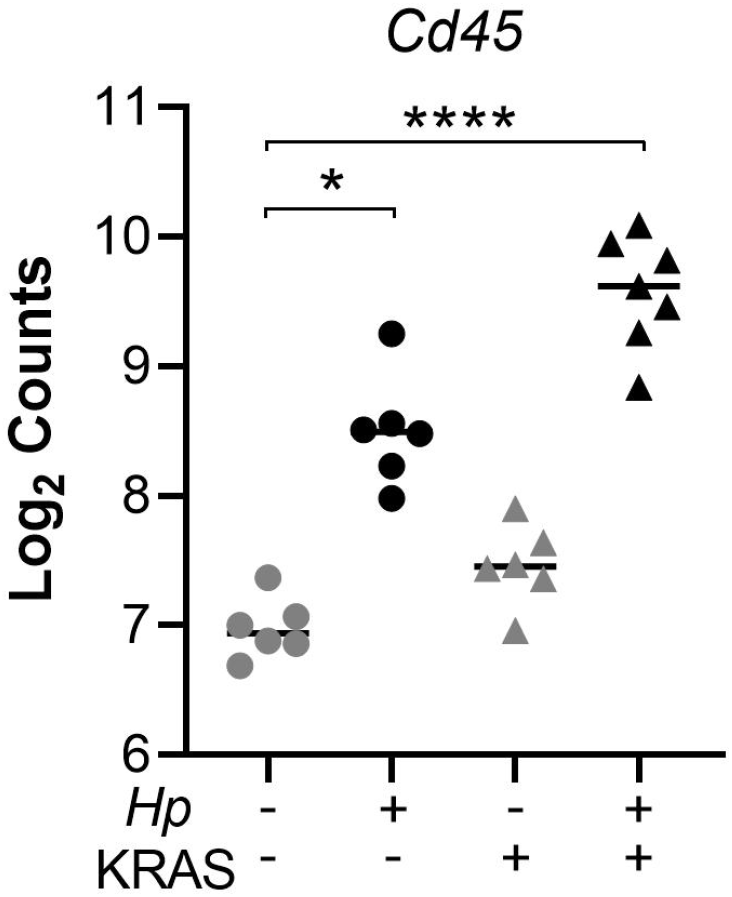
Infected mice have greater immune cell infiltration than mock-infected mice do at 12 weeks. RNA was extracted from stomach sections from *Hp*+/-, KRAS+/- mice at 12 weeks and immune-related gene expression was detected with the NanoString nCounter Mouse Immunology Panel. Expression of the pan-immune cell marker *Cd45* is shown. Data points represent actual values for each individual mouse and bars indicate median values. Statistically significant comparisons are indicated by: * *P* < 0.05, **** *P* < 0.0001, Kruskal-Wallis test with Dunn’s multiple test correction. Data comes from N=1 NanoString experiment with n=6-7 mice per group from N=2 independent mouse experiments.

**Figure S4.**
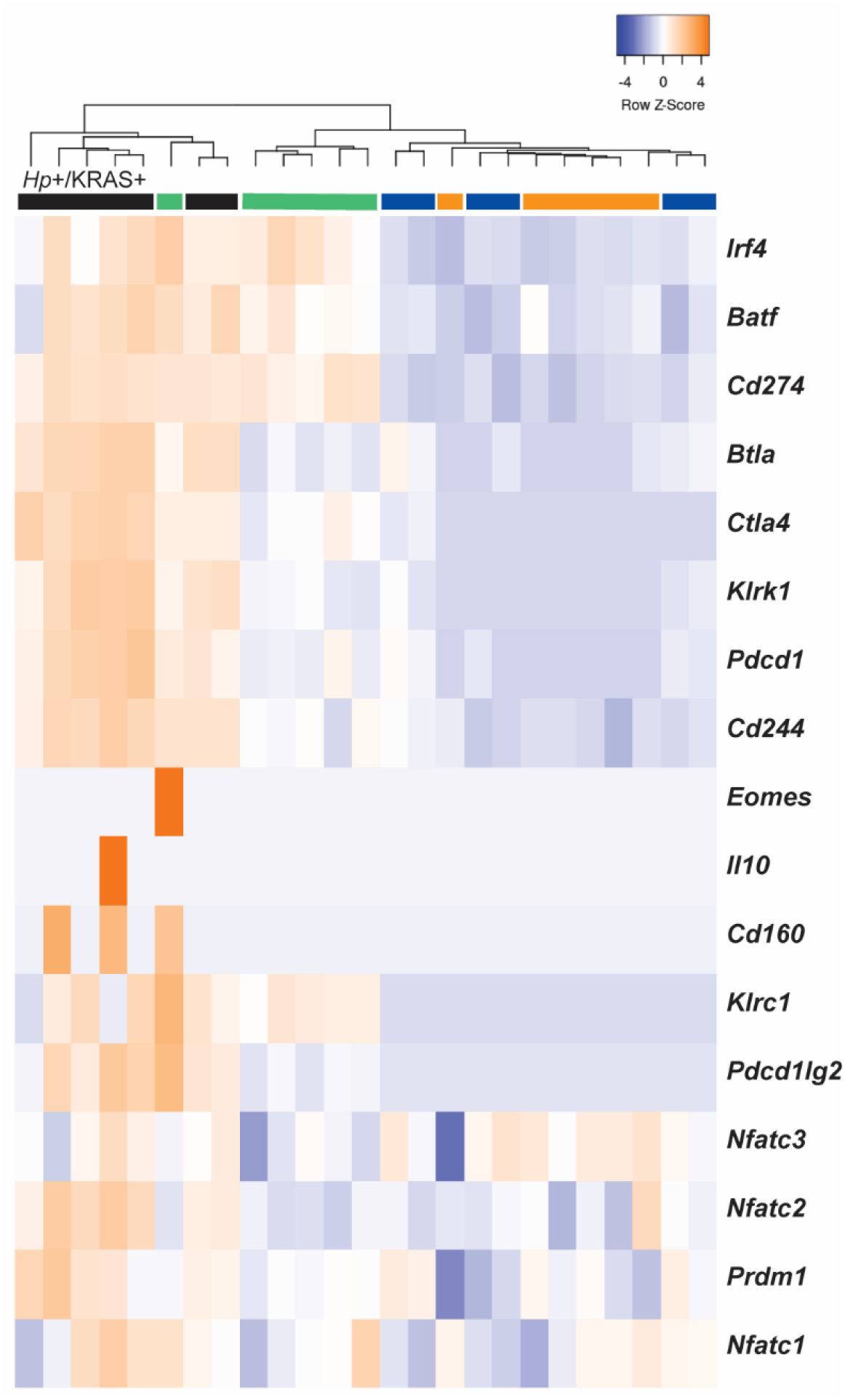
T cell exhaustion markers are upregulated in *Hp*+/KRAS+ mice at 12 weeks. RNA was extracted from stomach sections from *Hp*+/-, KRAS+/- mice at 12 weeks and immune-related gene expression was detected with the NanoString nCounter Mouse Immunology Panel. Expression of T cell exhaustion-related genes is shown. The dendrogram at the top of the heat map was produced by hierarchical clustering. Colored bars denote different treatment groups: orange is *Hp*-/KRAS-, green is *Hp*+/KRAS-, blue is *Hp*-/KRAS+, and black is *Hp*+/KRAS+. Data comes from N=1 NanoString experiment with n=6-7 mice per group from N=2 independent mouse experiments.

**Figure S5.**
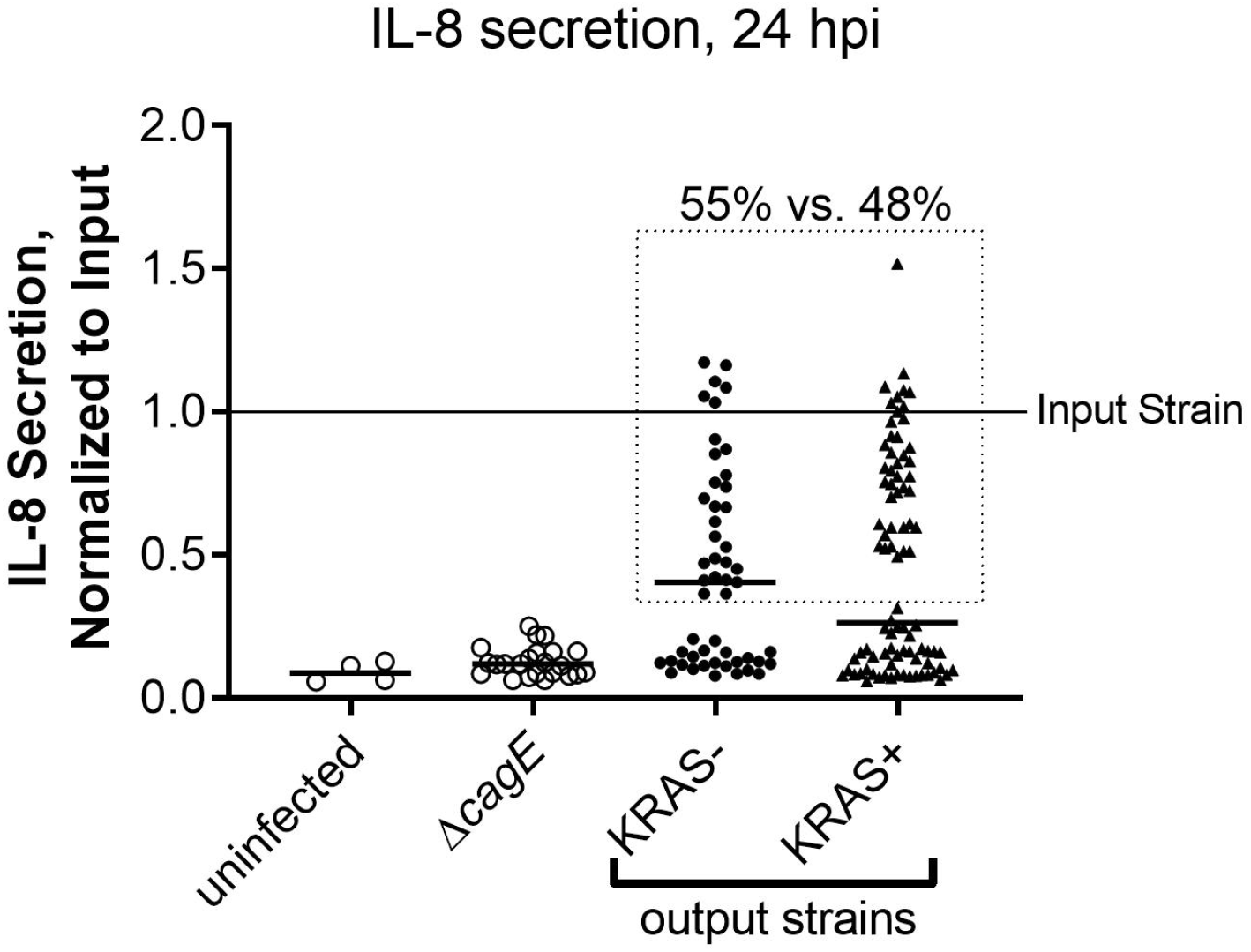
*Hp* isolates from KRAS- and KRAS+ mice exhibit similar retention vs. loss of type IV secretion system activity. The ability of *Hp* to elicit IL-8 secretion during co-culture with AGS gastric adenocarcinoma cells was used as a readout for functional type IV secretion. *Hp* strain PMSS1 was recovered from mice 12 to 15 weeks after active KRAS induction or sham induction (N=2 independent experiments), and individual colonies (n=5-6 per mouse) were used in triplicate co-culture experiments with AGS cells *in vitro*. At 24 hours post-infection, supernatants were collected and IL-8 was measured by enzyme-linked immunosorbent assay (ELISA). Absorbance values were normalized to the input strain (PMSS1 freezer stock). Shown are the average, normalized IL-8 values of individual *Hp* output isolates from KRAS- and KRAS+ mice. Uninfected AGS cells, and cells infected with PMSS1Δ*cagE*, which cannot assemble the type IV secretion system (T4SS), were included as controls. The difference in IL-8 secretion levels is not statistically significantly different (P > 0.05, Mann-Whitney U test). The dotted line encloses isolates deemed to have T4SS activity (55% of isolates from KRAS-mice and 48% of isolates from KRAS+ mice, P > 0.05, Fisher’s exact test).

**Figure S6.**
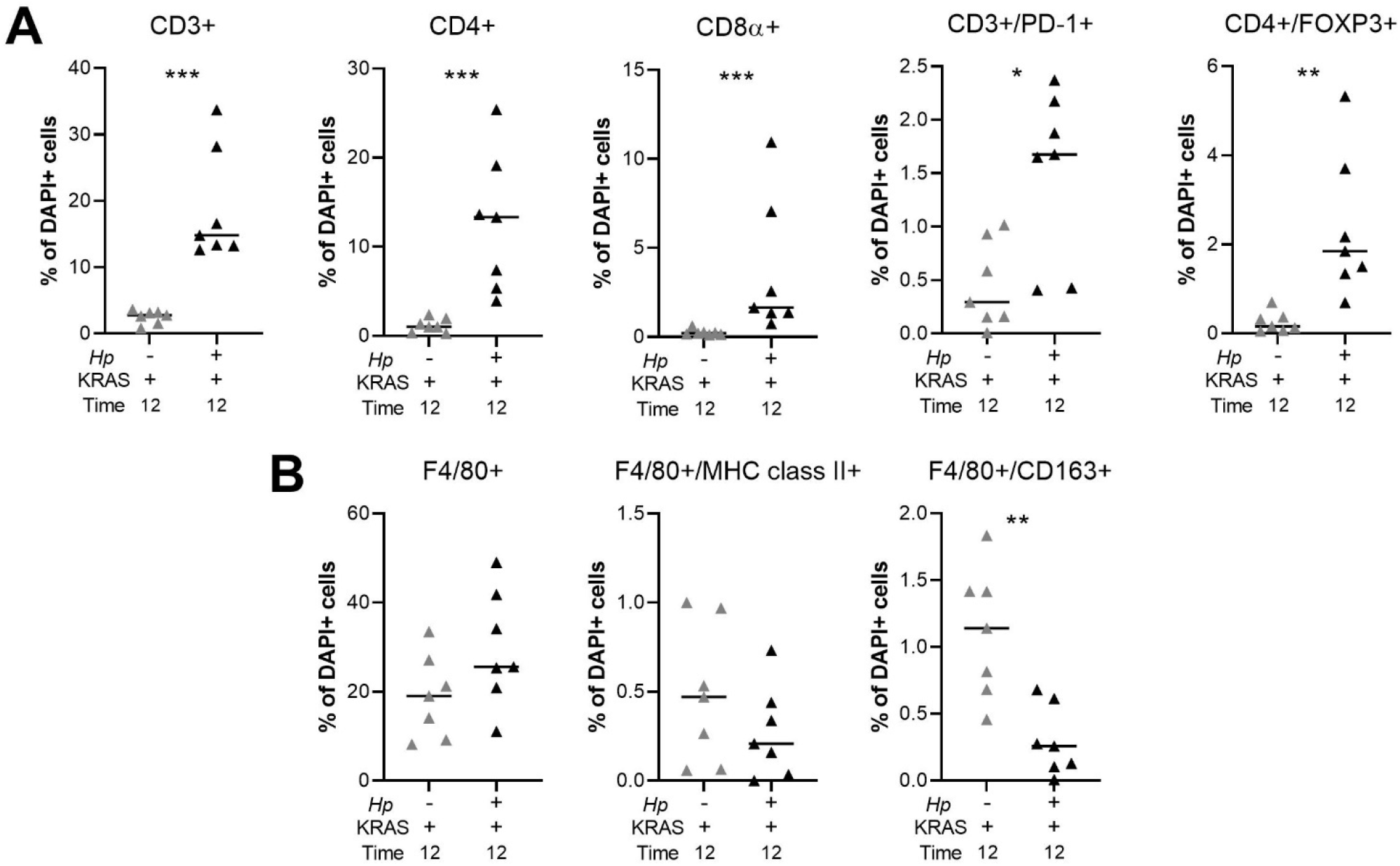
Quantitation of multiplex IHC confirms robust T cell infiltration and altered macrophage polarization in *Hp*+/KRAS+ mice. HALO software was used to detect and quantify the indicated T cell (**A**) and F4/80+ (**B**) subsets. Results are expressed as the percentage of all DAPI+ cells within a given image. Two images per mouse were detected and the average result for each marker is plotted. Bars indicate median values. Statistically significant comparisons are indicated by: * *P* < 0.05, ** *P* < 0.01, *** *P* < 0.001, Mann-Whitney U test.

**Figure S7.**
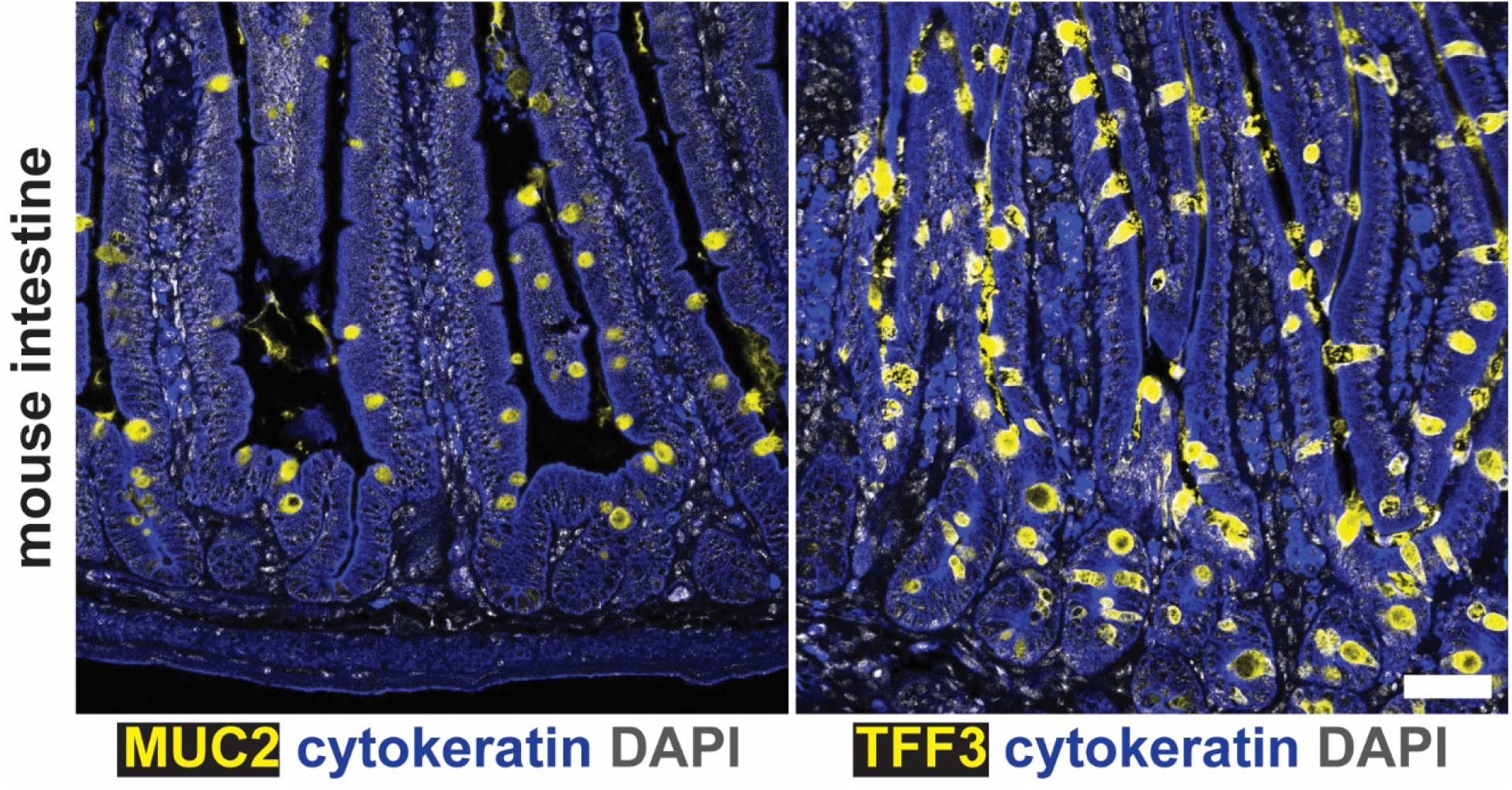
Goblet cells of mouse intestinal tissue stain positive for MUC2 and TFF3. Formalin-fixed, paraffin-embedded intestinal tissue from an untreated C57BL/6 mouse was deparaffinized and stained with the indicated antibodies in N=1 staining experiment conducted in parallel with the gastric staining shown in **Figure 6**. Representative images are shown. Scale bar, 50 µm.

**Figure S8.**
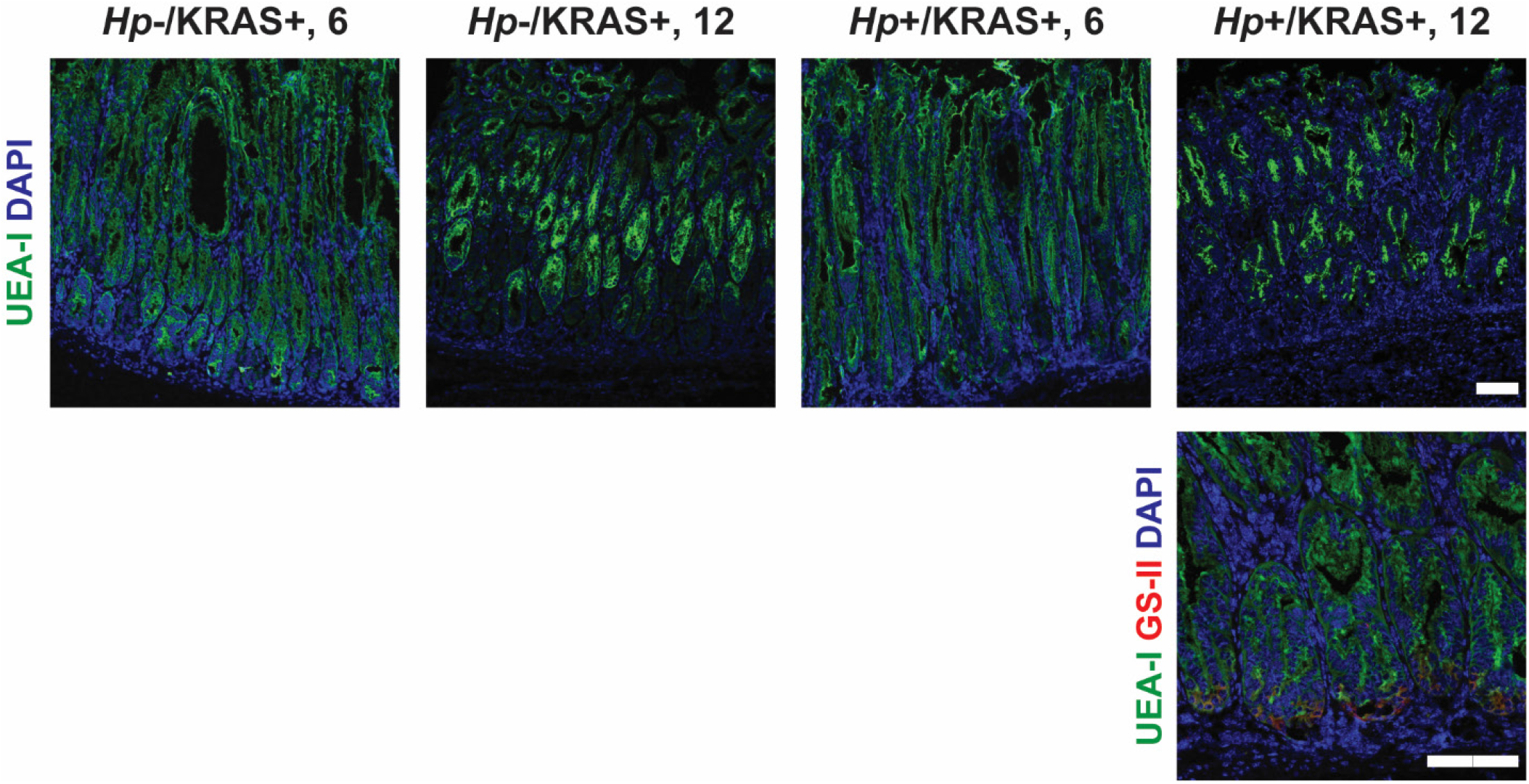
Foveolar hyperplasia does not differ between *Hp*-/KRAS+ and *Hp*+/KRAS+ mice. Corpus tissue from *Hp*-/KRAS+ and *Hp*+/KRAS+ mice obtained after six or 12 weeks (N=2 independent mouse experiments per time point) was assessed for foveolar hyperplasia via immunofluorescence microscopy. Stomachs were stained with the lectin UEA-I (green), which binds alpha-L-fucose in pit cells, as well as the lectin GS-II (red) and DAPI in N=2 staining experiments. Representative images are shown. Scale bars, 100 µm.

**Figure S9.**
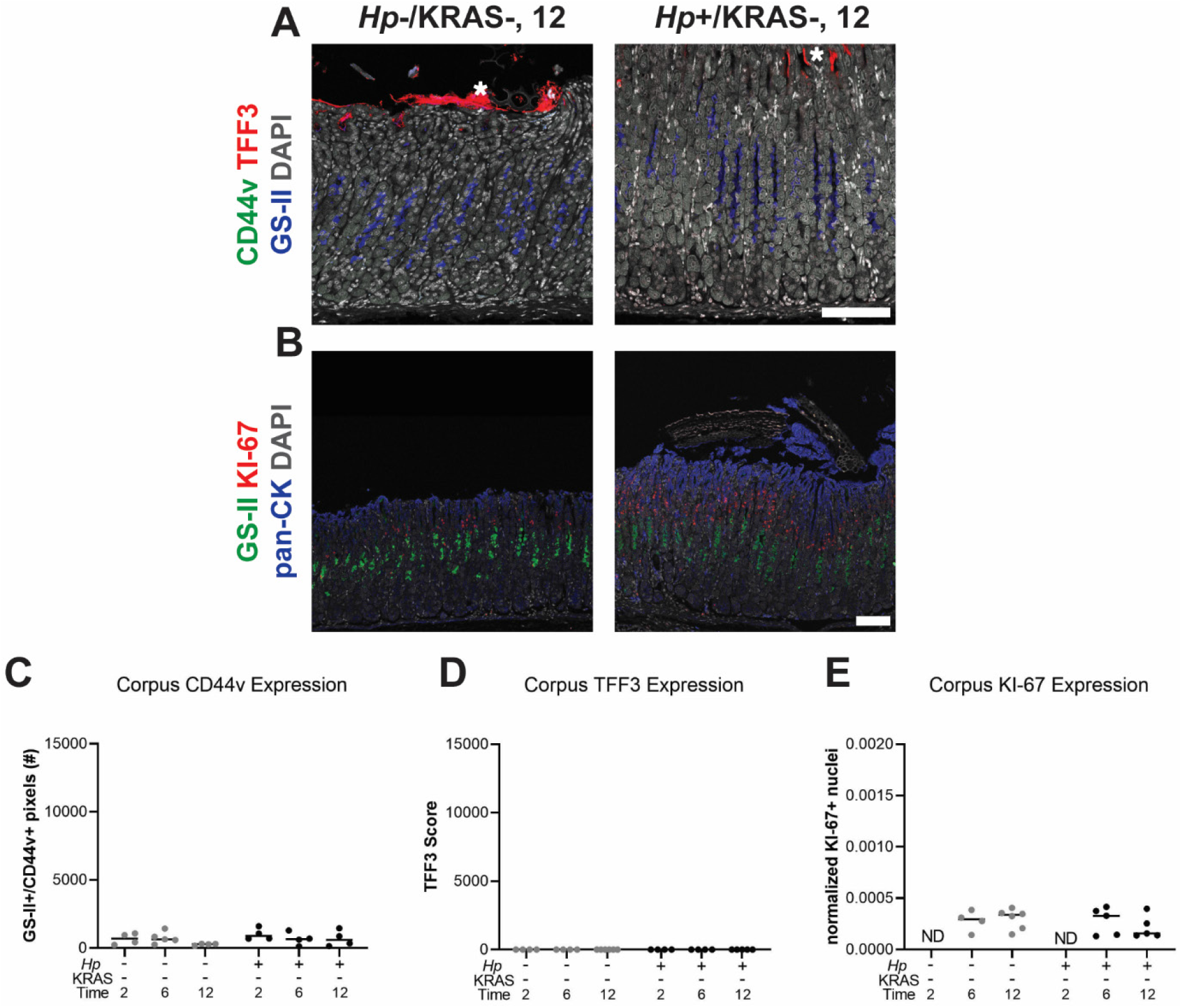
Metaplasia marker expression is not detected in KRAS-mice. Corpus tissue from *Hp*-/KRAS- and *Hp*+/KRAS-mice obtained after two, six or 12 weeks (N=2 independent mouse experiments per time point) was assessed for metaplasia via immunofluorescence microscopy. (**A and B**) (**A**) Stomachs were stained with antibodies against CD44v (green, no staining detected) and TFF3 (red), the lectin GS-II (blue) and DAPI (grey) in N=3 staining experiments. Asterisks show examples of non-specific (false-positive) TFF3 staining, which is not cell-associated, whereas specific staining is diffuse throughout the cytoplasm. (**B**) Stomachs were stained with antibodies against KI-67 (red) and pan-cytokeratin (blue), the lectin GS-II (green) and DAPI (grey) in N=3 staining experiments. Representative images are shown. Scale bars, 100 µm. (**C-E**) Three to five representative images per mouse were quantitatively or semi-quantitatively assessed and the median value for each mouse is plotted. Bars on the graphs indicate the median value for each mouse group. (**C**) The number of GS-II+/CD44v+ pixels per image was quantified. (**D**) TFF3 staining was semi-quantitatively scored in a blinded fashion. Non-specific staining was not included in the score. (**E**) KI-67+/DAPI+ nuclei were enumerated and normalized to the DAPI content (total number of DAPI+ pixels) of each image. ND, not determined.

**Figure S10.**
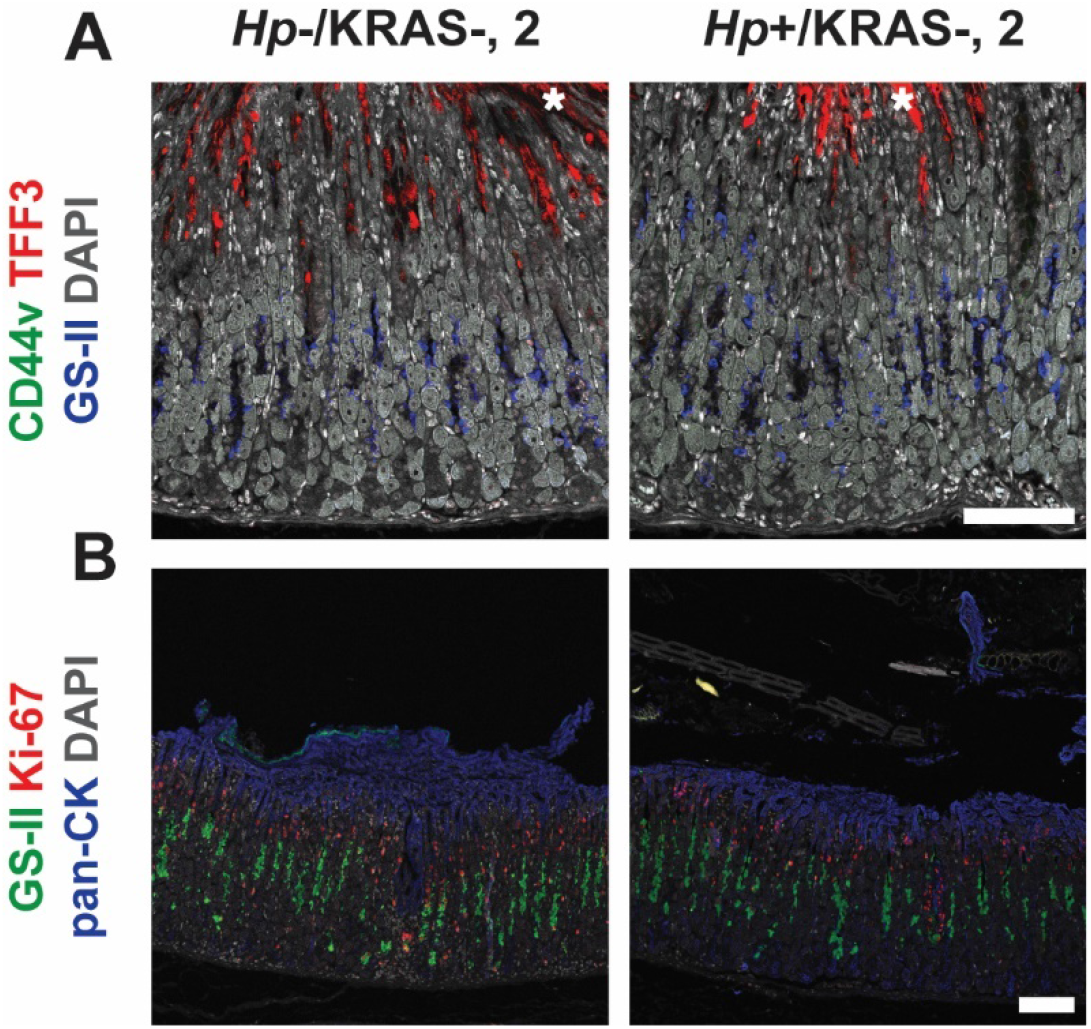
Metaplasia marker expression is low in KRAS+ mice at the two week time point. Corpus tissue from *Hp*-/KRAS+ and *Hp*+/KRAS+ mice obtained after two weeks (N=2 independent mouse experiments) was assessed for metaplasia via immunofluorescence microscopy. Representative images are shown. Scale bars, 100 µm. (**A**) Stomachs were stained with antibodies against CD44v (green, no staining detected) and TFF3 (red), the lectin GS-II (blue) and DAPI (grey) in N=3 staining experiments. Asterisks show examples of non-specific (false-positive) TFF3 staining, which is not cell-associated, whereas specific staining is diffuse throughout the cytoplasm. (**B**) Stomachs were stained with antibodies against KI-67 (red) and pan-cytokeratin (blue), the lectin GS-II (green) and DAPI (grey) in N=3 staining experiments.

**Figure S11.**
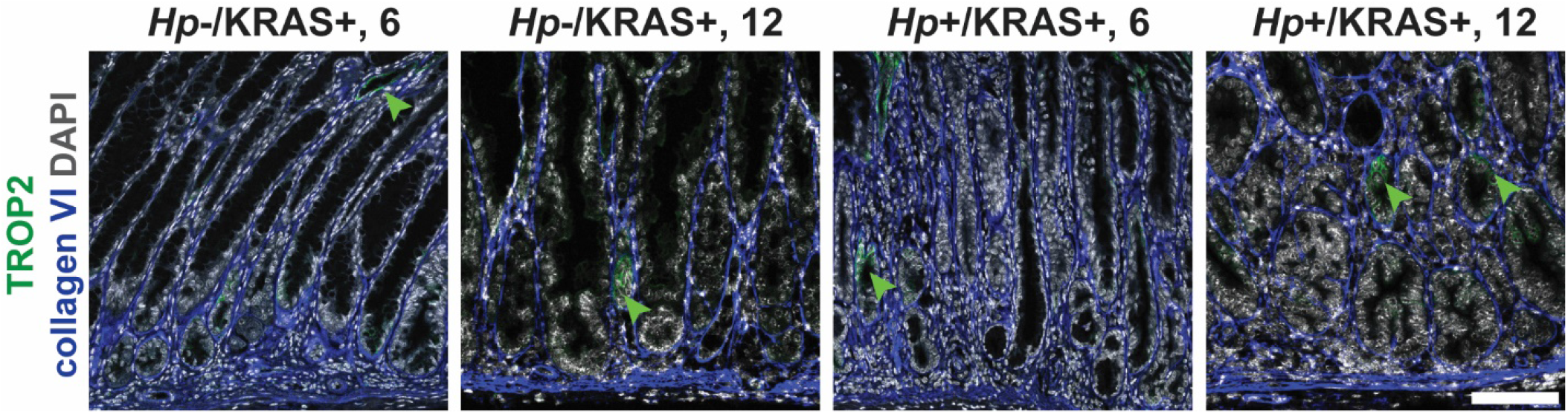
*Hp* infection increases TROP2 expression in KRAS+ mice. Stomachs from *Hp*-/KRAS+ and *Hp*+/KRAS+ mice obtained after six or 12 weeks (N=2 independent mouse experiments per time point) were assessed via immunofluorescence microscopy. Stomachs were stained with antibodies against TROP2 (green) and collagen VI (blue) and DAPI (grey) in N=3 experiments. Representative images are shown and arrows show TROP2+ glands. Scale bar, 100 µm. Collagen VI staining was used to detect individual gland fragments for TROP2 quantification.

**Figure S12.**
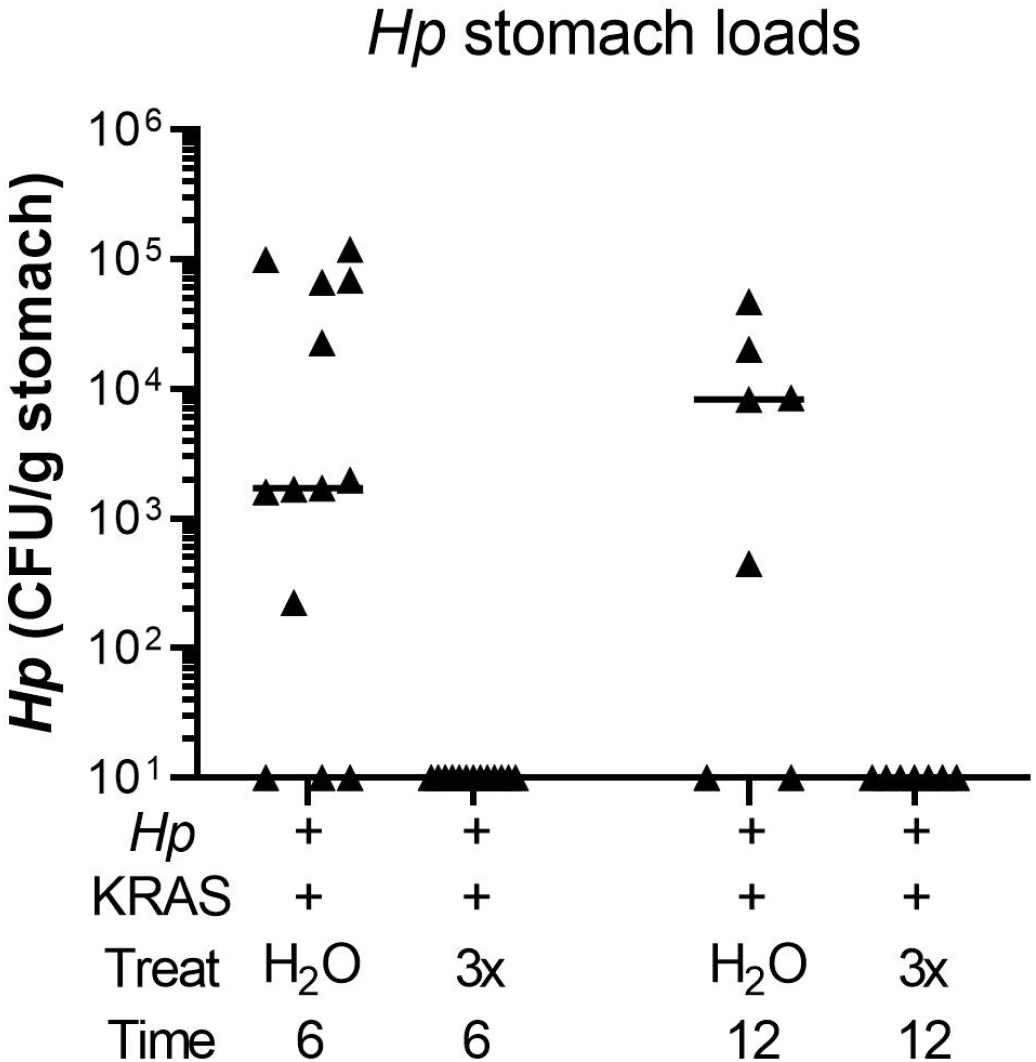
Antibiotic therapy eliminates *Hp* from the stomach. *Hp*+/KRAS+ mice received antibiotics (“3x”) or vehicle (“H_2_O”) after two weeks and were euthanized at six weeks (left), or received antibiotics or vehicle after six weeks and were euthanized at 12 weeks. *Hp* loads were assessed by quantitative culture; mice with no detectable colonization were plotted at the limit of detection.

**Supplementary Table 1.** Antibodies used to detect immune cell subsets via multiplex immunohistochemistry with the Leica BOND RX system.

| Position | Antibody | Clone & Host | Manufacturer & Catalog Number | Dilution (Conc.) | Secondary | Opal Dye |
| --- | --- | --- | --- | --- | --- | --- |
| 1 | MHC class II | M5/114.15.2 Rat | eBioscience 14-5321-85 | 1:200 (2.5 µg/ml) | ImmPress Rat-HRP | Opal 480 |
| 2 | PD-1 | D7D5W Rabbit | Cell Signaling #84651 | 1:50 (1.9 µg/ml) | Powervision Rabbit-HRP | Opal 650 |
| 3 | CD3 | SP7 Rabbit | Thermo RM-9107-S | 1:400 | Powervision Rabbit-HRP | Opal 520 |
| 4 | F4/80 | D2S9R Rabbit | Cell Signaling 20076S | 1:4000 (0.1 µg/ml) | Powervision Rabbit-HRP | Opal 540 |
| 5 | CD4 | 4SM95 Rat | eBioscience 14-9766-32 | 1:250 (2 µg/mL) | ImmPress Rat-HRP | Opal 570 |
| 6 | FoxP3 | FJK-16s Rat | eBioscience 14-5773-82 | 1:200 (2.5 µg/ml) | ImmPress Rat-HRP | Opal 620 |
| 7 | CD163 | EPR19518 Rabbit | Abcam ab182422 | 1:500 (1.4 µg/ml) | Powervision Rabbit-HRP | Opal 690 |
| 8 | CD8α | 4SM15 Rat | eBioscience 14-0808-82 | 1:1000 (0.5 µg/ml) | ImmPress Rat-HRP | Opal 780 |

**Supplementary Table 3.**
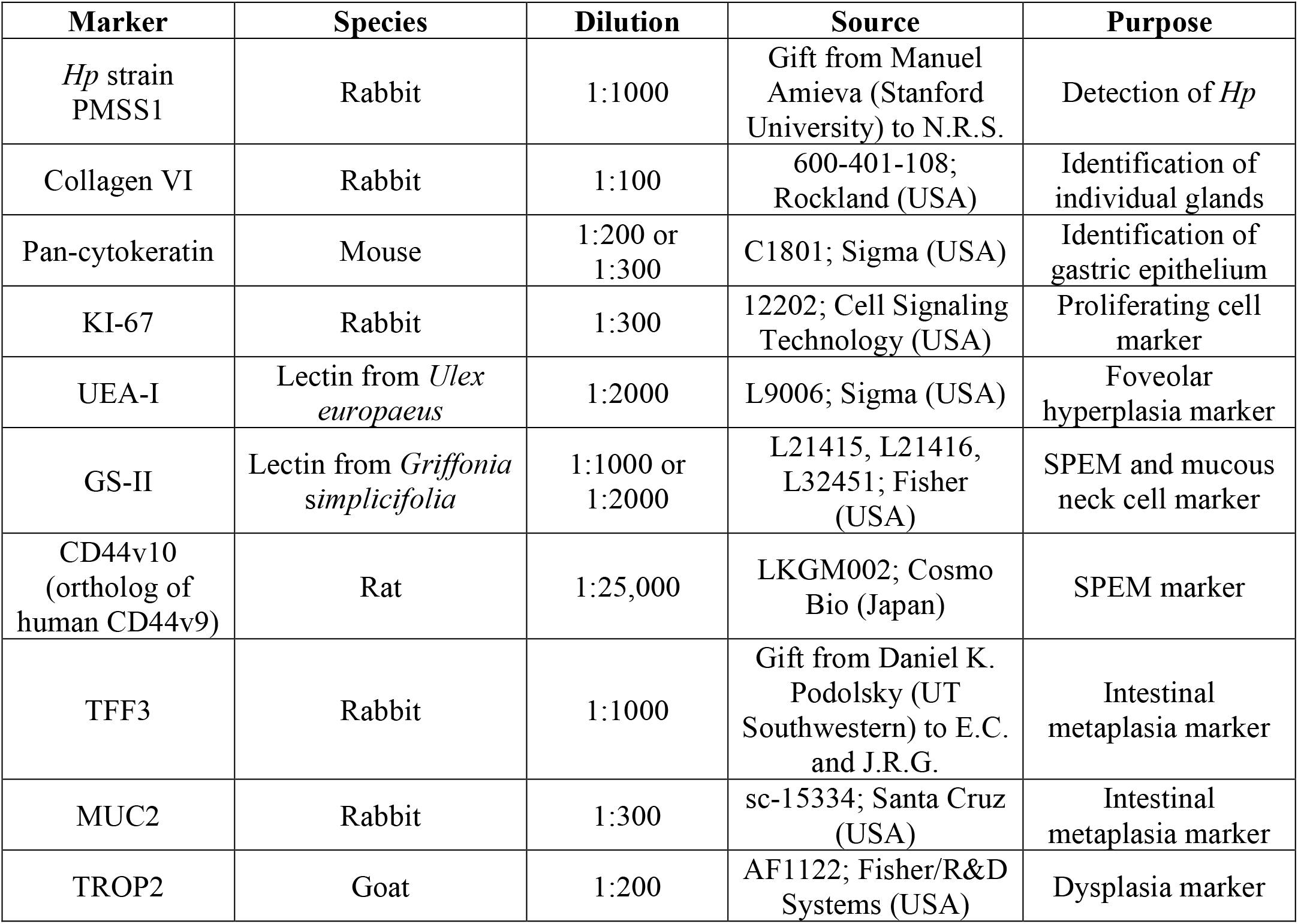
Antibodies and lectins used to assess metaplasia, dysplasia and cell proliferation in this study.

| Marker | Species | Dilution | Source | Purpose |
| --- | --- | --- | --- | --- |
| *Hp* strain PMSS1 | Rabbit | 1:1000 | Gift from Manuel Amieva (Stanford University) to N.R.S. | Detection of *Hp* |
| Collagen VI | Rabbit | 1:100 | 600-401-108; Rockland (USA) | Identification of individual glands |
| Pan-cytokeratin | Mouse | 1:200 or 1:300 | C1801; Sigma (USA) | Identification of gastric epithelium |
| KI-67 | Rabbit | 1:300 | 12202; Cell Signaling Technology (USA) | Proliferating cell marker |
| UEA-I | Lectin from *Ulex europaeus* | 1:2000 | L9006; Sigma (USA) | Foveolar hyperplasia marker |
| GS-II | Lectin from *Griffonia simplicifolia* | 1:1000 or 1:2000 | L21415, L21416, L32451; Fisher (USA) | SPEM and mucous neck cell marker |
| CD44v10 (ortholog of human CD44v9) | Rat | 1:25,000 | LKGM002; Cosmo Bio (Japan) | SPEM marker |
| TFF3 | Rabbit | 1:1000 | Gift from Daniel K. Podolsky (UT Southwestern) to E.C. and J.R.G. | Intestinal metaplasia marker |
| MUC2 | Rabbit | 1:300 | sc-15334; Santa Cruz (USA) | Intestinal metaplasia marker |
| TROP2 | Goat | 1:200 | AF1122; Fisher/R&D Systems (USA) | Dysplasia marker |

